# CCR2-targeting pepducins reduce T cell-nociceptor interaction driving bone cancer pain

**DOI:** 10.1101/2023.09.15.556569

**Authors:** Élora Midavaine, Rebecca L. Brouillette, Élizabeth Théberge, Christine E. Mona, Sakeen W. Kashem, Jérôme Côté, Vera Zeugin, Élie Besserer-Offroy, Jean-Michel Longpré, Éric Marsault, Philippe Sarret

## Abstract

Inhibition of the CCL2/CCR2 chemokine signaling represents a promising avenue for the development of non-opioid pain treatment, particularly for painful bone metastases. To investigate the involvement of CCR2 in cancer-induced bone pain, we generated and characterized the functional activities of a novel cell-penetrating pepducin, namely PP101, acting as an intracellular negative allosteric modulator of CCR2. *In vivo*, PP101 was effective in relieving neuropathic and bone cancer pain. By targeting CCR2, PP101 reduced bone cancer pain by preventing infiltration of CD4^+^ and CD8^+^ T cells and by decreasing the neuroimmune communication network within the dorsal root ganglia. Importantly, reduced neuroinflammatory milieu in the dorsal root ganglia induced by PP101 did not result in deleterious tumor progression or behavioral adverse effects. Thus, targeting the neuroimmune crosstalk through allosteric inhibition of CCR2 may represent an effective and safe avenue for the management of bone cancer pain.

**Graphical Abstract:** 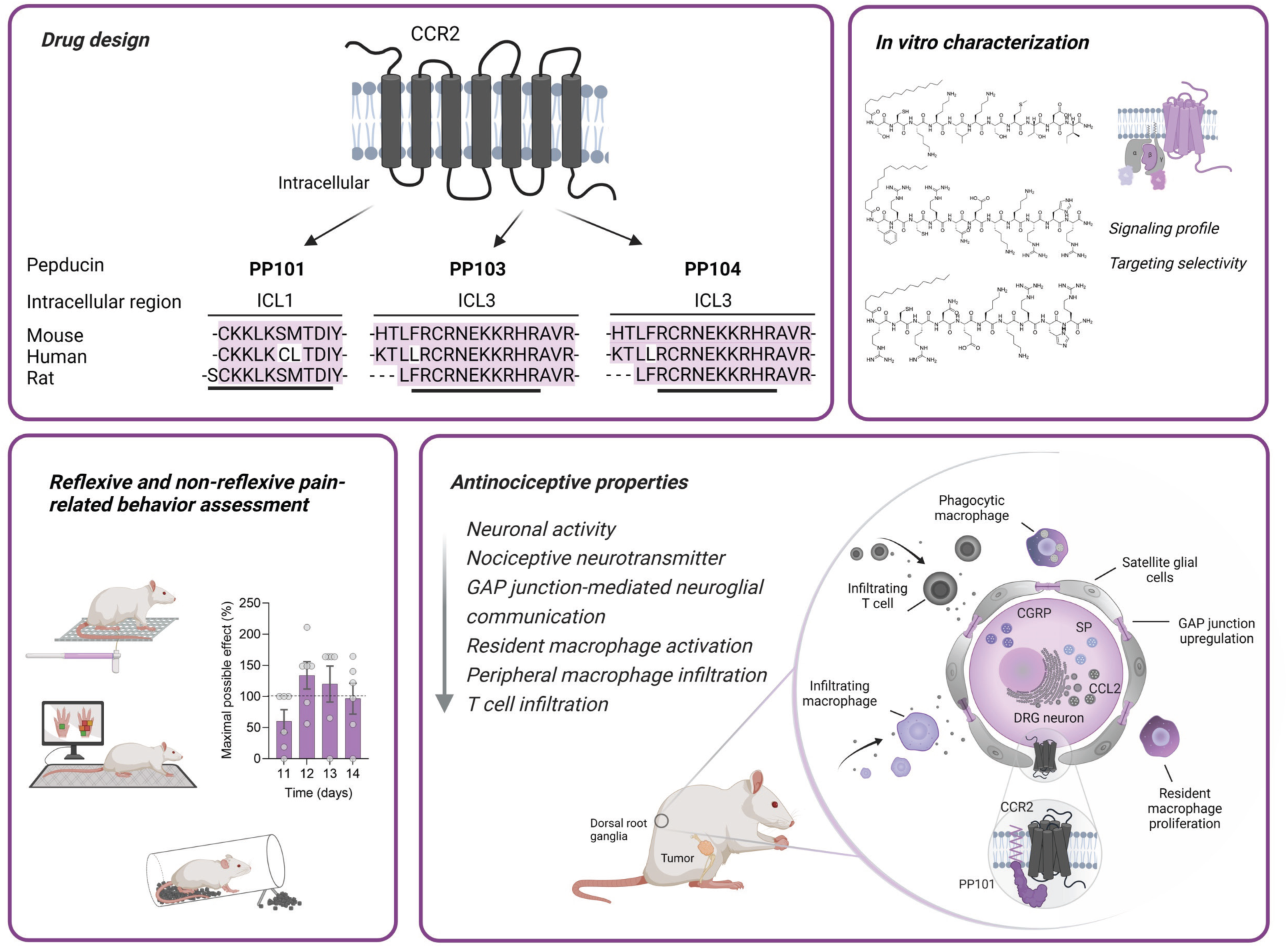

**Highlights:**

- Breast cancer bone metastases induce pain by activating CCR2 on sensory neurons.
- DRG-infiltrating CD4^+^ and CD8^+^ T cells promote the development of bone cancer pain.
- CCR2 inhibition by PP101 suppresses DRG neuroinflammation and neuronal excitability.
- PP101 alleviates bone cancer pain without behavioral or physiological side effects.

## Introduction

Bone cancer pain (BCP) is a chronic and debilitating condition, which affects more than 50% of women with metastatic breast cancer. Bone metastases cause severe pain with frequent breakthrough pain episodes for which the prescribed treatment regimens show very limited analgesic efficacy ^1^. The chemokine C-C ligand 2 (CCL2) and its cognate G protein-coupled receptor (GPCR) CCR2 have been shown to play key roles in nociceptive processing, mediating neuroinflammation, neuron-glia interaction and increase in the synaptic transmission in the spinal dorsal horn under chronic pain conditions ^2–4^. In addition, the CCL2/CCR2 axis was found to be overexpressed in various types of cancer with high metastatic potential, including aggressive triple-negative breast cancers ^5–7^. Importantly, elevated CCL2 levels are inversely correlated with patient survival, whereas CCR2 overexpression is positively correlated with clinical tumor stages^9–11^.

In recent years, the neuroimmune interface has received a growing interest in the field of pain research. Somatosensory neurons are indeed strongly modulated by alterations in glial and immune cell function ^12^. Likewise, spinal microglia and astrocytes actively regulate the neuronal network activity and are thus thought to act as central players in pain chronification ^13,14^. The spinal cord is an immune-privileged environment, protected by the blood-spinal cord barrier (BSCB). Neuroinflammatory diseases, such as multiple sclerosis (MS), experimental autoimmune encephalomyelitis (EAE) and neuropathic pain (NP), induce enhanced expression of inflammatory and chemotactic mediators, including cytokines and chemokines. As a result, this highly confined environment becomes more permeable to peripheral infiltrating blood-borne immune cells ^15–19^. On the other hand, dorsal root ganglia (DRG), lacking a blood-nerve barrier, are exposed to invading immune cells ^20^. To date, little is known about the involvement of resident and infiltrating immune cells in DRGs, and their respective contribution to cancer-induced bone pain.

GPCRs represent the main class of membrane receptors targeted by approved drugs ^21^. Given its important role in cancer biology and pain modulation, CCR2 appears as a promising druggable target. In this context, many orthosteric CCR2 antagonists have been developed and investigated with great success in preclinical models of diabetes, NP, rheumatoid arthritis, asthma, and MS, however, they all failed in clinical trials ^22^. The high redundancy and promiscuity in ligand-receptor interactions within the chemokine signaling network may explain the poor selectivity of CCR2 antagonists and therefore their lack of efficacy in clinical studies. In an attempt to overcome antagonists’ poor specificity, the anti-CCL2 antibody carlumab (CNTO 888) was developed and its efficacy examined in metastatic castration-resistant prostate cancer patients. Unfortunately, a 1000-fold increase in CCL2 serum concentration, in both patients and animals, raised safety concerns about the use of such medications ^23,24^. In addition, preclinical data demonstrated that interruption of anti-CCL2 antibody therapy leads to a dramatic breast cancer metastatic overshoot ^25,26^. Hence, targeting chemokine receptors at their orthosteric site yields many challenges that provide a rationale for alternative strategies.

In an effort to investigate the involvement of CCR2 in painful bone metastases, we developed and characterized cell-penetrating lipopeptide antagonists, called pepducins, targeting the intracellular domains of the receptor. We first assessed the analgesic potential of these CCR2-targeting pepducins in the breast MRMT-1 carcinoma model of cancer-induced bone pain in female rats. We also determined whether CCR2 antagonism induced changes in the molecular determinants of sensory neurons and in the recruitment of peripheral T cells into the DRGs. Finally, we examined if the development of BCP was associated with spinal glial reactivity, BSCB permeability and immune cell infiltration. Here, we report that BCP is mainly driven by molecular and cellular actors located in DRGs and that allosteric inhibition of CCR2 by pepducins is effective at reducing bone pain.

## Results

### Synthesis and characterization of a highly selective CCR2-inhibiting pepducin

In the present study, we developed pepducins, cell-penetrating lipopeptide antagonists of the chemokine receptor CCR2, to block the CCL2/CCR2 signaling. These lipopeptides were synthesized based on the first (ICL1) and third (ICL3) intracellular loops of the rat CCR2 receptor and coupled to an N-terminal palmitate moiety to facilitate their translocation across cell membranes as well as their attachment to the inner leaflet of the lipid bilayer (**Fig. 1A-C**). The high sequence homology of ICL1 and ICL3 between human, rat and mouse suggests that the CCR2 pepducins could possibly be active in all three species. Pepducins are lipopeptides that penetrate the cell by first anchoring their lipid moiety into the bilipid layer. The peptidic moiety then flips into the inner leaflet of the cell membrane, to interact with and modulate the target receptor. Because this penetration process is passive, dynamic, and bidirectional, the concentration of the peptide available to interact with the intracellular surface of the receptor is limited. As a consequence, the potency of pepducins for a specific receptor is reported to be 100 fold lower compared to classical extracellular orthosteric ligands that target the same receptor ^64^. All lipopeptides were first tested in BRET-based functional assay to assess their ability to modulate the canonical G_i/o_-protein and β-arrestin signaling pathways known to be activated by CCR2 agonists. In agonist mode, the ICL1- and ICL3-derived pepducins PP101, PP103, and PP104 were unable to activate Gα_i1_ and Gα_o_ proteins in HEK293 cells expressing CCR2, as opposed to CCL2 (**Fig. 1D and E**). In contrast, in antagonist mode, all three pepducins dose-dependently inhibited CCL2-induced Gα_i1_- and Gα_o_-protein activation, as well as β-arrestin 2 recruitment (**Fig. 1F-H**). Hence, these pepducins act as negative allosteric modulators of CCR2, with PP101 appearing to display the strongest inhibitory effect. Therefore, the selectivity of PP101 was further assessed on other class A GPCRs involved in nociceptive signaling and expressed in the DRG and spinal cord, such as the closely related chemokine receptors CXCR4 and CCR5, the µ-opioid receptor (MOR) and the neurotensin receptor type 1 (NTS1). Our results reveal that CXCR4-, CCR5-, MOR- and NTS1-mediated G-protein activation in response to SDF-1, CCL5, DAMGO and NT(8-13), respectively, were not antagonized by PP101 (**Fig. 1I-L, Supplementary Fig. 1A and B**). In addition, we found that the scrambled control pepducin SCR101, generated through randomly rearrangement of the original peptide sequence of PP101, was not able to antagonize CCL2-induced Gα_i/o_-protein activation (**Supplementary Fig. 1E-I**). Altogether, these results indicate that these pepducins act as negative allosteric modulators of CCR2 and that PP101 is selective for this receptor.

**Figure 1:**
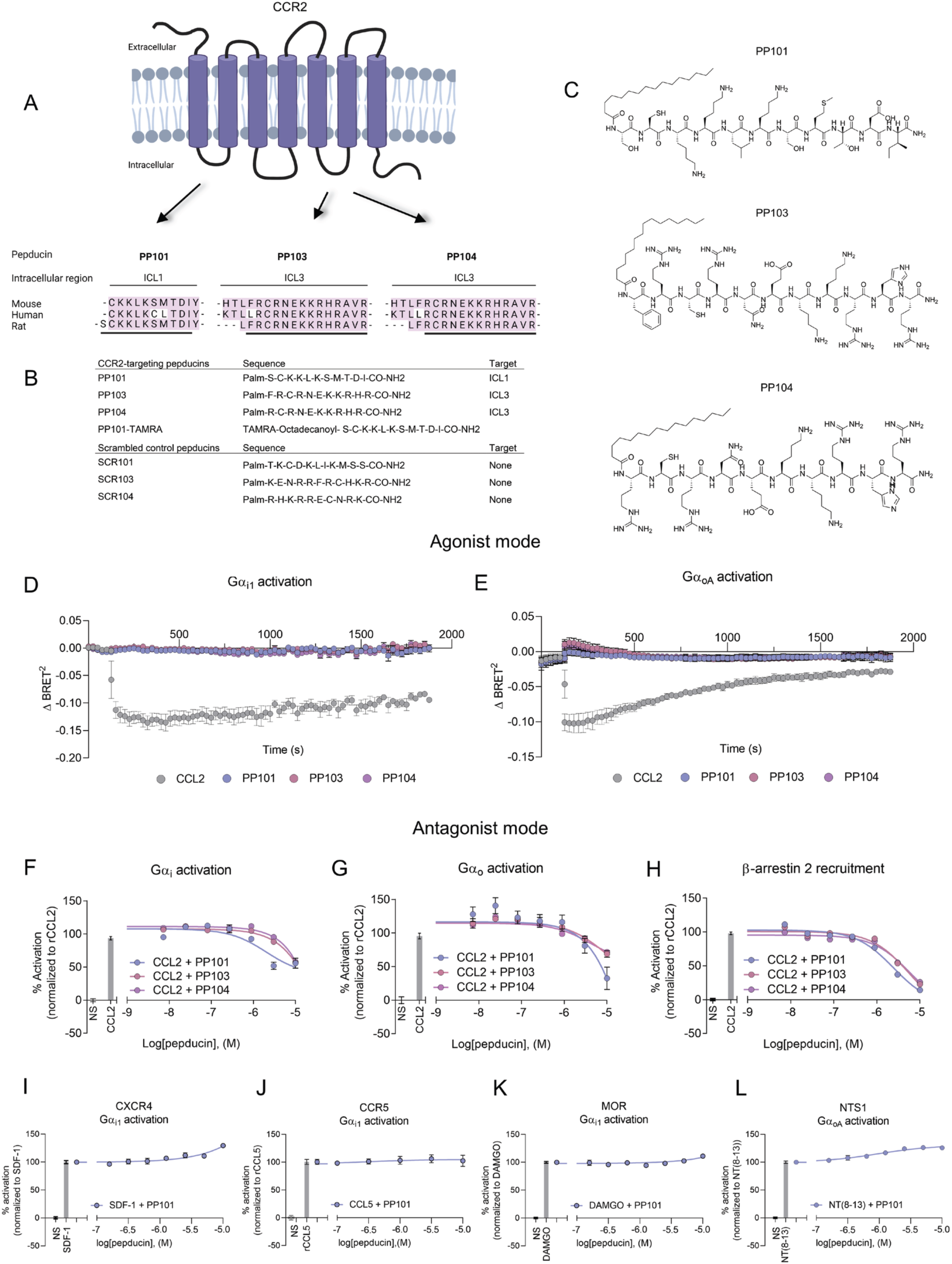
CCR2-targeting pepducins selectively inhibit CCL2-induced Gαi1/GαoA activation and β-arrestin2 recruitment through allosteric modulation. **(A)** Rational design of the pepducins derived from ICL1 (PP101) and ICL3 (PP103, PP104) of CCR2 and their respective amino acid sequences **(B-C)**. Time course of Gαi1 and GαoA activation upon CCL2 (2 µM) or pepducin (10 µM) stimulation assessed by measuring the BRET^2^ signal in real-time in HEK293 cells transiently transfected with rCCR2 and BRET^2^-based G-protein biosensors **(D-E)** (*n* = 5 duplicate). Antagonist effects of PP101, PP103 or PP104 on CCL2-induced Gαi1 or GαoA activation and β-arrestin 2 recruitment **(F-H)** (*n* = 3 duplicate). BRET^2^ ratios were normalized according to CCL2; values for HEK293 cells treated with CCL2 were set to 100% activation and the non-stimulated cells were set at 0%. Cross-antagonism of PP101 in G_i/o_-protein inhibition at CXCR4, MOR NTS1 or CCR5 stimulated respectively by SDF-1, DAMGO, NT(8-13) and rCCL5 were also assessed **(I-L)** (*n* = 3, quadruplicate). Palm: palmitate. Data are represented as mean ± SEM.

### PP101 reduces both stimulus-evoked and non-evoked bone cancer-related pain behaviors without deleterious adverse effects

Since the pepducin derived from the first intracellular loop of CCR2 exhibited the strongest antagonistic activity in *in vitro* assays (**Supplementary Fig. 2A-O**), PP101 was selected for further *in vivo* investigations. To assess the specificity of PP101 for CCR2, we first examined *in vivo* whether PP101 was able to reduce the mechanical hypersensitivity in a CCL2-induced acute pain model. As previously described ^27^, CCL2 acts as a neuromodulator in the spinal cord and its intrathecal (i.t.) administration in naive rats induces persistent mechanical allodynia (**Fig. 2A and B**). As opposed to rats receiving the inactive scrambled control pepducin SCR101, intrathecal delivery of PP101 was found to prevent the development of CCL2-induced acute mechanical hypersensitivity (**Fig. 2A and B**). Likewise, the selective CCR2 antagonist RS 504393 was effective at blocking CCL2-evoked allodynia, which supports that the antiallodynic effect of PP101 was mediated by the inhibition of CCR2.

**Figure 2:**
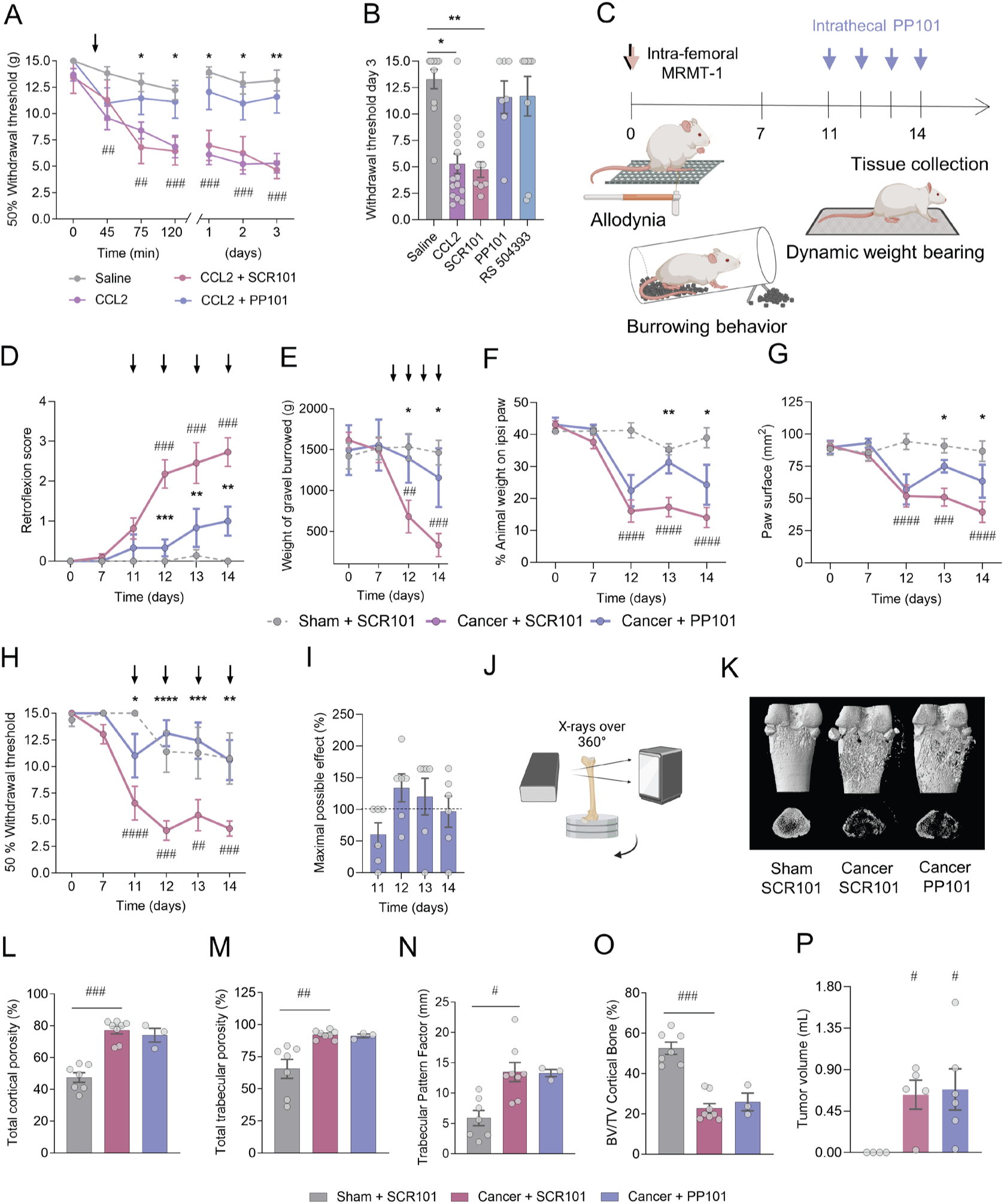
The pepducin PP101 prevents the nociceptive behaviors induced by bone cancer without deleterious effect on bone remodeling and tumor growth. (**A**) Effect of CCL2 (1 µg/rat, i.t.) and co-injection of CCL2 with PP101 (125 nmol/rat, i.t.) on mechanical thresholds, compared to rats receiving SCR101 control pepducins. (**B**) Paw withdrawal thresholds measured at day 3 following i.t. saline (*n* = 14), CCL2 (*n* = 6), CCL2+SCR101 (*n* = 7), CCL2+PP101 (*n* = 8) and CCL2+RS 504393, a selective CCR2 antagonist (60 nmol/rat; *n* = 9). (**C**) Schematic representation of the study design and of the behavioral tests used to measure the stimulus-evoked and non-evoked pain-related behaviors in tumor-bearing female rats. Repeated daily administrations of PP101 or SCR101 (125 nmol/rat, i.t.) between days 11 and 14 on (**D**) paw retroflexion score, (**E**) burrowing behavior, (**F and G**) dynamic weight-bearing and (**H-I**) mechanical allodynia, (sham+SCR101 *n* =7, cancer+SCR101 *n* = 7, cancer+PP101 *n* = 6). (**J-O**) Tomodensitometry (µCT) analysis of the bone structure of sham-operated and tumor-implanted femur (sham+SCR101 *n* = 7, cancer+SCR101 *n* = 8, cancer+PP101 *n* = 6). (**P**) Volume of the tumor on the ipsilateral femur (sham+SCR101 *n* = 4, cancer+SCR101 *n* = 5, cancer+PP101 *n* = 8). Data are represented as mean ± SEM. Two-way ANOVA followed by Sidak’s multiple comparisons test in (**A, D-H**), Kruskal-Wallis followed by Dunn’s multiple comparisons test in (**B, L-P**). \**P* < 0.05, \*\**P* < 0.01, \*\*\**P* < 0.001. # compared with sham+SCR101 and * compared with cancer+SCR101.

The analgesic effectiveness of the pepducin PP101 was then monitored in a model of cancer-induced bone pain which consists in implanting breast MRMT-1 tumor cells into the femoral medullary cavity of female Sprague-Dawley rats. To this end, rats received daily doses of PP101 (125 nmol/rat; intrathecal) from postoperative days (POD) 11 to 14 (**Fig. 3C**). Our results reveal that PP101 was able to significantly reduce bone tumor-induced paw retroflexion, a coping behavior developed to limit paw use and contact with the floor during rest and ambulation (**Fig. 3D, Supplementary Fig. 3A-E**). Using non-reflexive behavioral assays as proxies of global well-being and functional disability, we further demonstrated that PP101 treatment induced a maintenance of spontaneous burrowing behaviors as well as increased the weight born and paw surface applied on the cancer-bearing paw in freely moving rats (**Fig. 2E-G**). In addition, the pepducins targeting CCR2 reversed the development of sensory-reflexive behaviors compared to SCR101-treated tumor-bearing littermates in response to von Frey filament stimulations, reaching maximal efficacy (**Fig. 2H and I**). Since cancer-induced bone pain has a neuropathic component ^1^, we further assessed the ability of PP101 to reverse the nociceptive behaviors associated to neuropathic pain. Our results demonstrate that repeated intrathecal administrations of PP101 attenuated the mechanical hypersensitivity in female rats that underwent chronic constriction injury (CCI) of the sciatic nerve (**Supplementary Fig. 3F**). Importantly, CCR2-targeting pepducins did not alter animal’s daily body weight gain (**Supplementary Fig. 3G**), pain thresholds, or induce side effects in naive animals (**Supplementary Fig. 4A-L**), nor did they induce toxicity in cultured DRG neurons (**Supplementary Fig. 4M-O**), thus indicating that repeated treatment with PP101 is safe. Finally, we investigated whether negative allosteric modulation of CCR2 by chronic intrathecal administrations modulated tumor-induced bone resorption. As determined by post-mortem microcomputed tomodensitometry (μCT) imaging analyses, sustained PP101 i.t. treatment did not alter tumor-induced bone loss (**Fig. 2J and K**). Indeed, all morphological parameters, including cortical and trabecular bone porosity, trabecular bone pattern factor and cortical bone volume ratio (BV/TV) were similar to SCR101-treated tumor-bearing rats (**Fig. 2L-O**). Moreover, the tumor volume also remained unaltered following PP101 treatment, suggesting that the analgesic effects of PP101 after i.t. administrations were restricted to central actions (**Fig. 2P**). Moreover, this suggests PP101 was devoid of deleterious effect on bone remodeling and tumor progression which is expected after intrathecal administrations. Together, these results demonstrate that CCR2-targeting pepducins are effective in reducing the reflexive and non-reflexive pain-related behaviors caused by bone tumor growth and that the analgesic action of PP101 is devoid of behavioral or physiological side effects.

**Figure 3:**
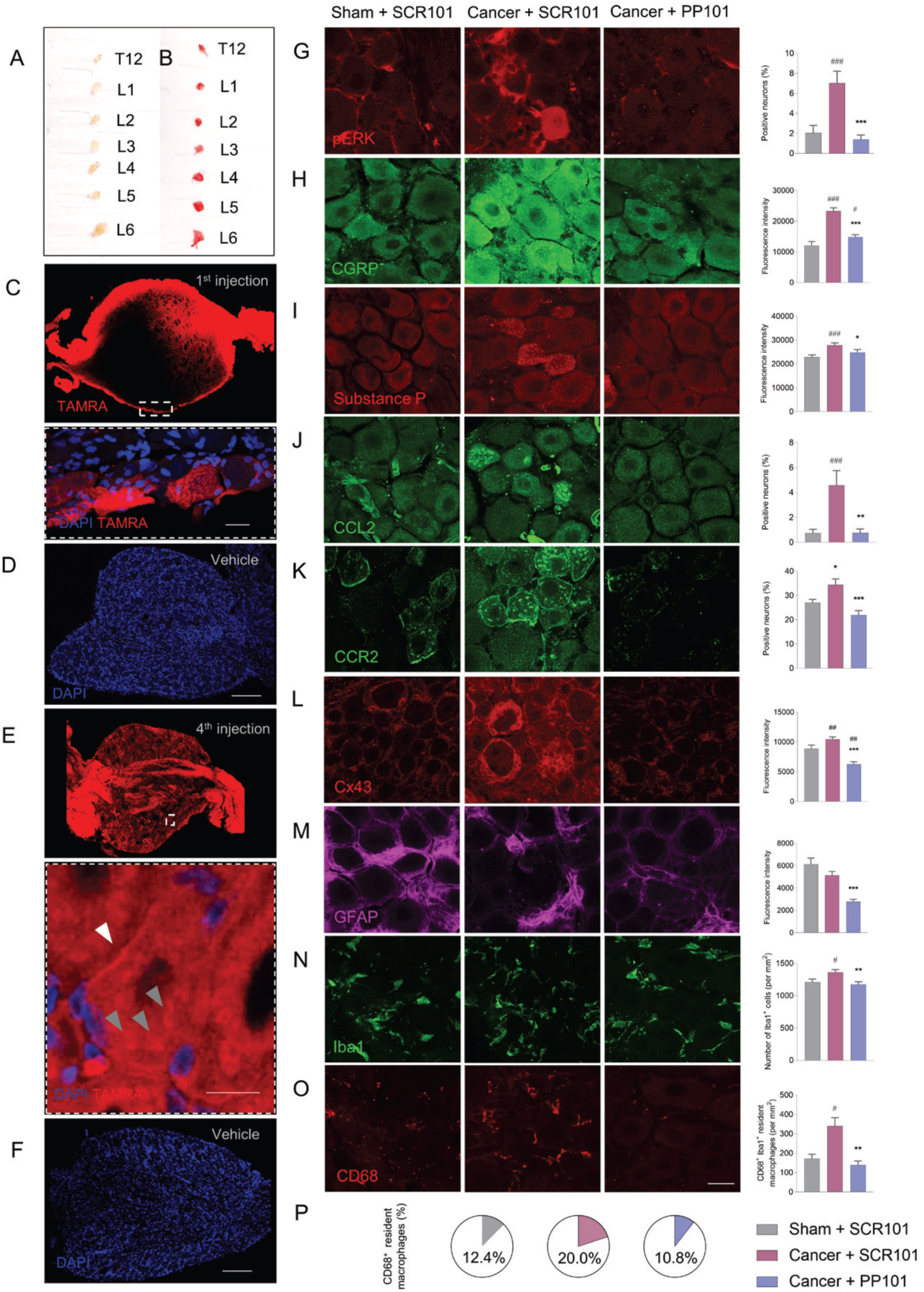
CCR2 allosteric antagonism attenuates the neuronal phenotypic changes induced by bone cancer pain in the DRGs. (**A-D**) Detection of PP101-TAMRA in DRGs, 45 min after the first injection and its accumulation throughout the DRG following four injections (**E-F**). PP101-TAMRA is found in neuronal membrane (white arrowhead) and intracellular vesicles (gray arrowheads) (*n* = 3 rats per group, all lumbar DRG were analyzed, L3 is shown in (**C-F**). Scale bars in **C**, **E**, **E (insert**) and **F** correspond to 20 µm, 150 µm, 10 µm and 150 µm, respectively. Effect of PP101 on (**G**) ERK activation, (**H**) CGRP, (**I**) substance P, (**J**) CCL2, (**K**) CCR2, (**L**) connexin 43 (Cx43) expressed by satellite glial cells (SGC), (**M**) GFAP expression by SGC, (**N**) Iba1^+^ resident macrophages and (**O**) CD68^+^ phenotype at POD 14. (**P**) Percent of CD68^+^ macrophages in the Iba1^+^ macrophage population in experimental groups. (*n* = 3 animals per group, 8-10 slices of L1-3 DRG analyzed per animal). Data represented as mean ± SEM. Kruskal-Wallis followed by Dunn’s multiple comparisons test in (**G-O**). Scale bar corresponds to 20 µm. \**P* < 0.05, \*\**P* < 0.01, \*\*\**P* < 0.001. # compared with sham+SCR101 and * compared with cancer+SCR101.

### Allosteric inhibition of CCR2 alters the cancer-induced neuroinflammatory landscape of the dorsal root ganglion

Bone cancer pain depicts a unique and poorly understood neurochemical signature compared to other types of chronic pain, such as inflammatory and neuropathic pain ^1^. Hence, we sought to identify the underlying mechanisms associated with the analgesic action of PP101 by focusing on the primary sensory neurons of lumbar DRGs which relay peripheral nociceptive signals to the spinal cord and exhibit increased activity in chronic pain conditions. First, we examined the tissue distribution of the ICL1-derived pepducin into DRGs following i.t. administration, by linking a TAMRA fluorophore to the N-terminal end of PP101 (PP101-TAMRA) (**Supplementary Fig. 1C, D and I**). Macroscopic visualization confirmed that PP101-TAMRA effectively penetrates lumbar DRGs (**Fig. 3A and B**). 45 min post-injection, confocal examination under higher magnification revealed the presence of PP101-TAMRA in sensory neurons located around the circumferential border of the DRG and penetrating about half of the DRG parenchyma (**Fig. 3C and D**). Importantly, PP101-TAMRA was found to accumulate and distribute throughout the DRG in response to the four-injection paradigm, used here to determine the *in vivo* functional activity of PP101 (**Fig. 3E and F**).

We next investigated whether repeated administration of PP101 was able to downregulate the neurochemical changes accompanying the neuronal activation during bone cancer pain. At POD 14, the enhanced neuronal activity was readily detectable by the increased number of pERK-positive neurons, as well as by the increased immunoreactivities of CGRP and substance P (**Fig. 3G-I, Supplementary Fig. 7H and I**). Accordingly, expression of the neuronal injury marker ATF3 was also induced in different subpopulations of sensory neurons in tumor-bearing rats (**Supplementary Fig. 3H and I**). Likewise, we observed an increased percentage of CCL2- and CCR2-positive neurons compared to sham animals, supporting the key role of the CCL2/CCR2 axis in driving neuroinflammation and bone cancer pain (**Fig. 3J and K, Supplementary Fig. 7J**). On the other hand, there was an increase in gap junction-forming protein connexin 43 (Cx43), revealing extensive paracrine communications between satellite glial cells (SGC) and sensory neurons during the development of bone cancer pain (**Fig. 3L**). Tumor-induced bone pain also induced expansion and activation of Iba1^+^ resident macrophages with no effect on GFAP immunoreactivity in surrounding satellite glial cells (**Fig. 3M-P**). Importantly, our results demonstrate that CCR2 allosteric antagonism with PP101 significantly decreased ERK activation, CCL2/CCR2 chemokine signaling, and pain-related neuropeptide expression in sensory neurons (**Fig. 3G-K**). In addition, it reduced the expansion and phagocytic phenotype of resident macrophages (**Fig. 3M-P**) as well as the immunoreactivity of Cx43 in SGC of cancer-bearing rats (**Fig. 3L**). Taken together, these data demonstrate that the CCR2-derived pepducins were effective at attenuating the changes to both neuronal and glial phenotypes which contribute to the excitation of sensory neurons, the loss of neuronal shielding and the potentiation of the glial-neuronal crosstalk underlying the development of bone cancer pain.

### PP101 blocks bone cancer pain by inhibiting T cell infiltration into dorsal root ganglia

Immune cells are known as noteworthy contributors to pain signaling. Accordingly, targeted therapies against peripheral immune cell infiltration into the peripheral and central nervous system were shown effective at alleviating neuropathic pain ^28^. To gain further insight into the involvement of infiltrating immune cells in cancer-induced bone pain, we assessed the inhibitory potential of PP101 on their recruitment and phenotype into the affected DRGs. Importantly, bone cancer pain stimulated the recruitment of peripheral immune cells into the DRG displaying the signature markers of T cells (CD3^+^) and monocytes (CD11b) (**Fig. 4A-B, Supplementary Fig. 7G, K and M**). Furthermore, CCR2^+^ immune cells were found in, and restricted to, the leptomeninges of tumor-bearing female rats, as they are absent from the DRG parenchyma (**Supplementary Fig. 6A-B**). In addition, T cells were found to infiltrate the satellite glial cell sheath and to be in the vicinity of neuronal somata (**Supplementary Fig. 6C**), as previously described following nerve damage ^29^. Repeated treatment with PP101 was found to decrease the infiltration of CD3^+^ T cells and monocytes in DRGs as well as CCR2^+^ immune cells in leptomeninges (**Fig. 4A-B, Supplementary Fig. 6A-B**). Importantly, the analgesic effect of repeated PP101 administrations was associated with a decrease in both CD4^+^ and CD8^+^ T cells while the phenotype of infiltrating macrophages remained unaltered (**Fig. 4C-G, Supplementary Fig. 5A-H**).

**Figure 4:**
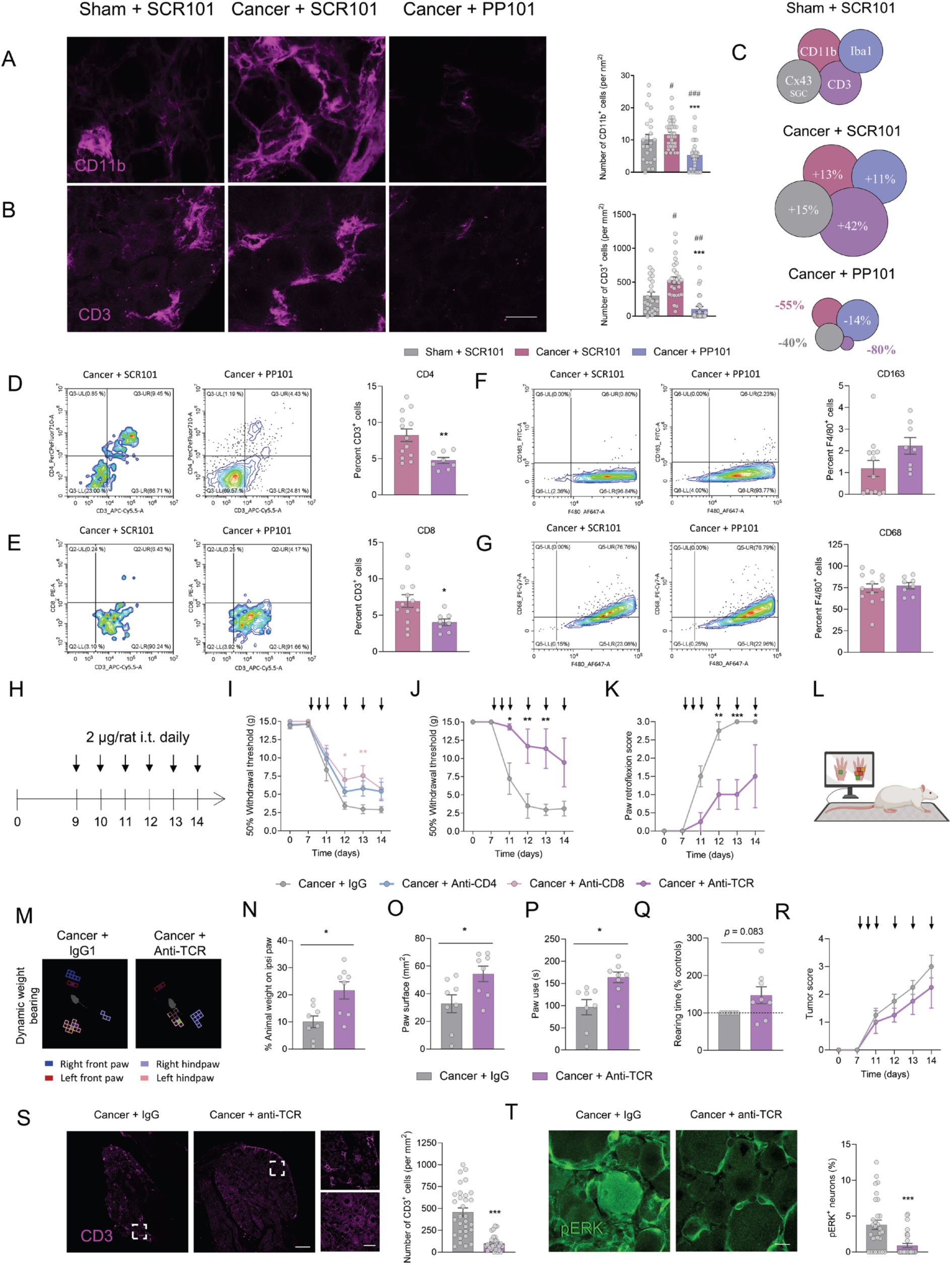
T-cell infiltration into the dorsal root ganglion contribute to bone cancer pain and is reduced by PP101. Effect of the CCR2-targeting pepducin, PP101 on (**A**) CD11b^+^-infiltrating macrophages and (**B**) CD3^+^ T-cells at POD 14 (*n* = 3 per group, 8-10 slices of L1-3 DRG analyzed per animal). (**C**) Schematic representation of the dynamic of Cx43^+^ satellite glial cells (SGC), CD11b^+^, Iba1^+^ and CD3^+^ populations in DRG tissue of sham and cancer tumor-bearing rats receiving SCR101 or tumor-bearing rats treated with PP101. Percent of (**D**) CD4^+^ and (**E**) CD8^+^ infiltrating CD3^+^ T cells analyzed by flow cytometry. Percent of (**F**) CD68^+^ and (**G**) CD163^+^ infiltrating F4/80^+^ monocytes analyzed by flow cytometry (cancer+SCR101 *n* = 13, cancer+PP101 *n* = 8). (**H**) Repeated daily administrations of anti-CD4, anti-CD8 or anti-IgG control antibody (2 µg/rat, i.t.) between days 9 and 14 on mechanical allodynia. Repeated daily administrations of anti-TCRαβ or anti-IgG control antibody (2 µg/rat, i.t.) between days 9 and 14 on (**I, J**) mechanical allodynia and (**K**) paw retroflexion score in tumor-bearing animals. (**L**) Schematic representation of the dynamic weight bearing device and (**M**) representative weight acquisition for anti-TCRαβ- and anti-IgG antibody-injected tumor-bearing rats. (**N-Q**) Dynamic weight-bearing measurements performed at POD 13 and 14, (**R**) tumor score, and histological analysis of (**S**) CD3^+^ T-cells and (**T**) pERK^+^ neurons in anti-TCRαβ and anti-IgG antibody-injected tumor-implanted animals at POD 14 (cancer+anti-IgG *n* = 4, cancer+anti-TCRαβ *n* = 5). Data represented as mean ± SEM. Kruskal-Wallis followed by Dunn’s multiple comparisons test in (**A and B**), Mann-Whitney test in (**D-G, N-Q, and S-T**), Two-Way ANOVA followed by Sidak’s multiple comparison test in (**I-K and R**). Scale bar corresponds to 20 µm. \**P* < 0.05, \*\**P* < 0.01, \*\*\**P* < 0.001. # compared with sham+SCR101 and * compared with cancer+SCR101 or cancer+IgG1.

Since T lymphocytes were the cell population with the greatest increase in DRG induced by bone tumor, and the cell population with the largest decrease after chronic administrations of PP101 (**Fig. 4C**), the involvement of T cells in cancer-induced bone pain was selectively investigated by depleting T cell subsets in DRGs. Chronic i.t. administration of anti-CD8 antibodies partially reversed bone cancer pain whereas anti-CD4 antibodies lacked significant analgesic effects (**Fig. 4H and I**). In contrast, administration of anti-TCRαβ antibodies, targeting both CD4 and CD8 subsets, to tumor-bearing animals prevented the development of mechanical allodynia and weight bearing-deficits compared with controls receiving anti-IgG injections (**Fig. 4J-Q**). Despite the known role of T cells in cancer surveillance, PP101 and neutralizing antibodies were here administered into the intrathecal space, avoiding any possible modulation of the tumor growth localized in the femur bone. Indeed, these compounds are likely contained within the injection site due to the presence of the meningeal layers. Thus, the pain behaviors reduced by specific T-cell depletion were independent of an effect on tumor growth and led to a reduction in neuronal activation, as determined by reduced neuronal p-ERK immunoreactivity (**Fig. 4R-T**). Thus, peripheral immune cells act as drivers of DRG neuron activation and bone cancer-induced hypersensitivity and inhibition of T cell infiltration by PP101 blocks bone cancer pain.

### Bone cancer pain is independent of microglial activation or spinal immune cell infiltration

We next explored the contribution of the neuroimmune signaling within the spinal nociceptive circuits of tumor-bearing female rats. The neuron-specific nuclear protein NeuN revealed a slight but significant increase in ERK MAPK signaling into the spinal cord marginal (lamina I) neurons 14 days after cancer cell inoculation, as compared to sham animals (**Fig. 5A and B**). Using immunohistochemistry, we further demonstrated that bone cancer also promoted faint but significant c-fos immediate-early gene induction in nociceptive neurons located in the superficial dorsal horn (**Fig. 5C**). Chronic intrathecal delivery of PP101 was found to reduce the spinal lamina I neuronal activity, as shown by the decrease in pERK and c-fos immunoreactivity (**Fig. 5B and C**).

**Figure 5:**
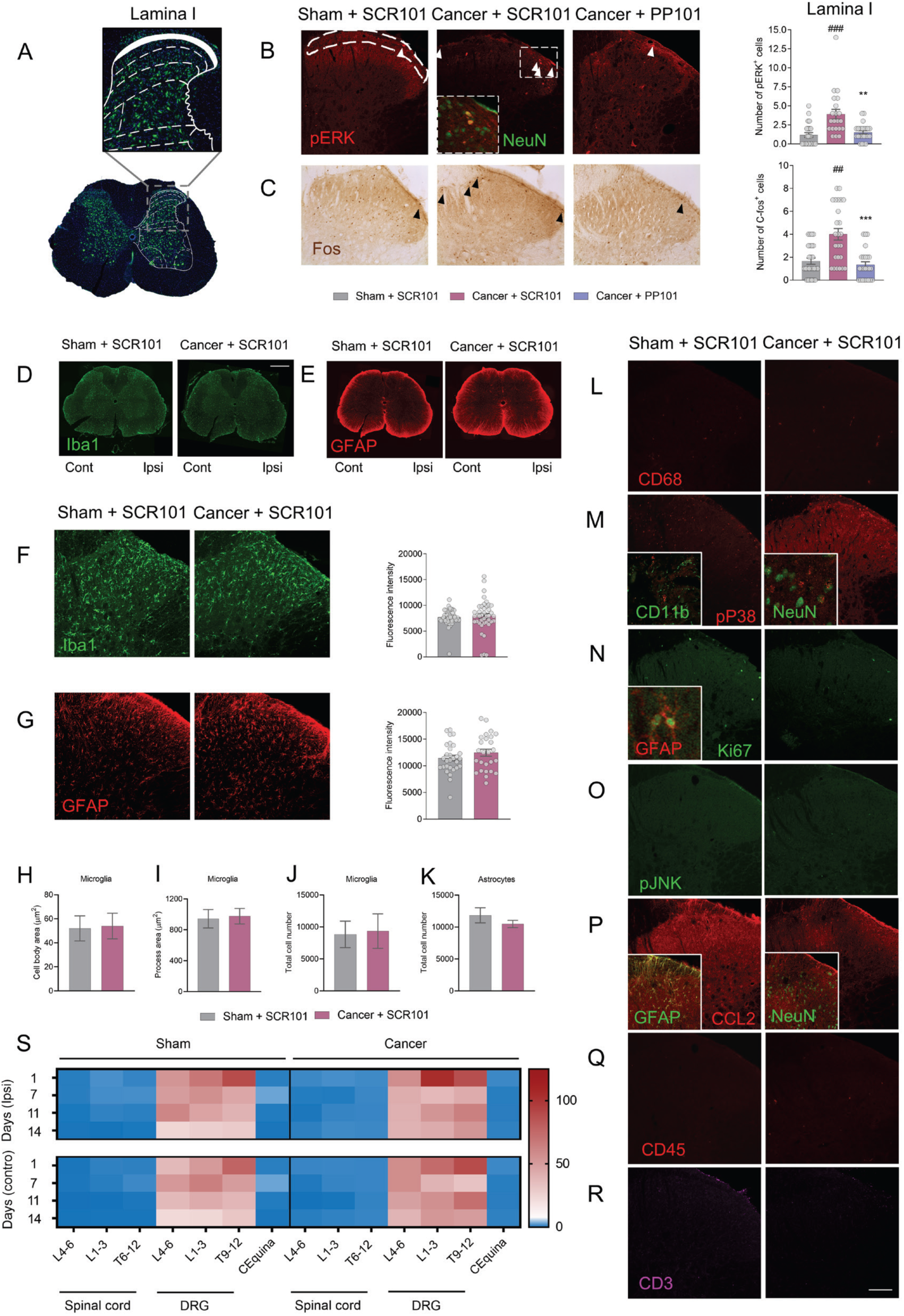
Bone cancer pain induces lamina I spinal neuron activation, which is reduced by PP101 but does not require glial cell activation or immune cell infiltration into the spinal cord. (**A**) Representation of the nociceptive-specific spinal lamina I. Effect of PP101 on (**B**) pERK and (**C**) C-Fos activation in lamina I neurons (white and black arrowheads, respectively). (**D-E**) Whole slide imaging of Iba1 and GFAP glial cell markers. Scale bar corresponds to 500 µm. (**F**) Iba1-positive microglia and (**G**) GFAP-reactive astrocytes in the ipsilateral superficial dorsal horn at POD 14 in tumor-bearing female rats. (**H-K**) Surface area of the microglial soma and processes and total cell number of microglia and astrocytes (*n* = 3 per group analyzed in triplicate). (**L-N**) Lack of CD68-, phospho-p38 MAPK- and Ki67-expressing microglia as well as absence of pJNK MAPK- and CCL2-stained astrocytes (**O-P**). Absence of staining for the CD45 pan-immune cell and CD3 T-cell specific markers in the spinal dorsal horn (**Q-R**) (*n* = 3 per group, 10 slices of L2-3 spinal cord analyzed per animal). Scale bar corresponds to 100 µm. **(S)** Spinal cord, DRG and *cauda equina* permeability to sodium-fluorescein at POD 1, 7, 11 and 14 (*n* = 3 per group analyzed in triplicate). Data are represented in µg/g tissue. Data represented as mean ± SEM. Mann-Whitney test in (**B-C**), Kruskal-Wallis followed by Dunn’s multiple comparisons test in (**F-K**). \**P* < 0.05, \*\**P* < 0.01, \*\*\**P* < 0.001. # compared with sham+SCR101 and * compared with cancer+SCR101.

The development of chronic pain has also been associated to spinal neuroinflammatory responses, as characterized by glial cell reactivity, peripheral immune cell infiltration and BSCB permeability^30^. In tumor-bearing female rats, analysis of the spinal glial cell populations at POD 14 revealed a striking absence of activated microglia (Iba1) and reactive astrocytes (GFAP) (**Fig. 5D-G**). Accordingly, no changes in the morphology and density of Iba1-labeled microglia or in the total cell number of astrocytes were observed (**Fig. 5H-K**). In various chronic pain models, microglial cells were shown to exhibit increased p38 MAPK signaling to develop a phagocytic phenotype (CD68^+^) and to locally proliferate (Ki67^+^) around highly active nociceptive neurons. However, no increase in phospho-p38 MAPK, CD68 or Ki67 immunoreactivity could be observed over the microglial population at POD 14 (**Fig. 5L-N**). pP38 MAPK colocalized with neurons (NeuN) but not with microglial cells (CD11b) in cancer-bearing animals. Rare Ki67^+^ cells could be observed in the astrocytic population but not in microglia. At the molecular level, astrocytes showed no sign of activation, as revealed by the absence of activation of JNK MAPK signaling and decreased in CCL2 immunolabeling, previously identified as markers of astrocyte-mediated pain persistence (**Fig. 5O and P**). The absence of CD3^+^ T cells and CD45^+^ immune cells in the spinal dorsal horn also indicated that the infiltration of peripheral immune cells into the spinal cord is unlikely to be involved in the development of bone pain in female rats, in contrast to DRGs (**Fig. 5Q and R; Supplementary Fig. 7F-G and L-M**). We finally investigated the permeability of the BSCB at different time points (POD 1, 7, 11 and 14) following tumor cell inoculation. Along with the lack of immune cell infiltration into the spinal cord, the BSCB remained intact at all time points studied (**Fig. 5S**). Together, these data show limited contribution of spinal glial cells or spinal cord infiltrating peripheral immune cells in tumor-induced bone pain in female rats.

## Discussion

Chemokines play a key role in the initiation and maintenance of persistent pain states and are therefore potential therapeutic targets for pain management ^2–4^. However, the promiscuity and redundancy of the chemokine system pose considerable challenges for the development of drugs targeting specific chemokine receptors and may explain the failure of chemokine receptor antagonists in clinical trials ^31^. Here, we thus explored the therapeutic potential of cell-penetrating lipopeptides targeting CCR2 and acting as negative allosteric modulator to relieve bone cancer pain. We first demonstrated *in vitro* the selectivity of PP101 for the CCR2 receptor over class A GPCRs involved in pain signaling and expressed in DRGs. We also assessed the selectivity of PP101 *in vivo* by its ability to block CCL2-induced acute hypersensitivity. We then demonstrate that PP101 was effective in reducing bone tumor-induced reflexive and non-reflexive pain-related behaviors. Allosteric inhibition of CCR2 significantly reduced neuronal expression of the CCL2/CCR2 system as well as pain-related neuropeptides SP and CGRP in primary afferent neurons. The downregulation of CCR2 was unexpected, but most likely results from a decrease in neuronal activity, as CCR2 is upregulated following neuronal excitability in chronic pain conditions ^32^.

A key concept emerging from our findings is that nociceptor-T cell interactions in the DRG have a critical role in bone cancer pain. Accumulating evidence indicates that T cells contribute to the transition from acute to chronic pain, but the precise localization of this neuron-adaptative immune cell crosstalk remains controversial ^33,34^. Our results show that activation of primary sensory neurons releases the pronociceptive and chemotactic neurotransmitters SP, CGRP, and CCL2, likely contributing to T cell infiltration into DRGs ^35–37^. Importantly, chronic PP101 treatment in tumor-bearing animals reduced infiltration of CD4^+^and CD8^+^ T cells subsets into the DRG parenchyma, as well as reflexive and non-reflexive pain behaviors. Accordingly, we found that sustained neutralization of TCRαβ^+^ T cells, but not CD4^+^ or CD8^+^ subsets alone, in the DRG compartment, completely blocked the development of bone cancer pain. Despite the growing evidence for the role of infiltrating T cells in the development of neuropathic pain ^33,34^, this is, to our best knowledge, the first demonstration of the contribution of infiltrating T cells in the DRG in bone cancer pain. In addition, the expression of the gap-junction protein connexin 43 is up-regulated in DRG satellite glial cells of tumor-bearing animals. This loss of the neuronal shielding, which is thought to play an important role in maintaining peripheral sensitization, favors neuronal crosstalk and paracrine signaling between neuronal cell bodies, therefore opening the gate for synchronized firing among ensemble of neurons ^38^. CCR2 antagonism by PP101 also decreased the expression of Cx43 in satellite glial cells surrounding the neuron cell bodies, thus providing preservation of neural somata shielding. Aside from the inflammatory component of bone cancer pain, we should not exclude the possibility that the analgesic effect of CCR2-targeting pepducins may also result from the inhibition of pain induced by bone tumor-induced nerve entrapment and compression, as PP101 also reverses nociceptive behaviors in a model of nerve constriction injury.

Furthermore, our results reveal limited contribution of spinal cord infiltrating peripheral immune cells to the maintenance of bone cancer pain. Increasing evidence suggests that peripheral immune cells invading the spinal cord play an important role in the chronic pain pathophysiology, especially in neuropathic pain conditions ^17,39–43^. Accordingly, an increased permeability of the BSCB correlates with the enhanced capacity of these T lymphocytes to migrate toward the spinal cord ^15,16^. However, some reports have recently questioned the involvement of spinal cord-infiltrated T cells to the development of neuropathic pain ^44,45^. Unlike suggested previously in a mouse model of bone cancer pain in male ^18^, we show here that the spinal cord parenchyma of tumor-bearing rats is devoid of infiltrating T cells and monocytes at POD14, which is consistent with the maintenance of the BSCB integrity. This apparent discrepancy between both studies may be due to sex differences in the biological mechanisms underlying the development of bone cancer pain ^46–49^.

There is an abundant literature supporting the role of resident glial cells such as microglia and astrocytes within the spinal cord in the pathogenesis of chronic pain ^50,51^. However, while glial cell activation in the spinal cord seems to be a prerequisite step for pain chronification in male, the involvement of spinal glial cells to the pathobiology of neuropathic pain in female is currently a subject of controversy ^46,52–54^. As previously suggested ^55–57^, we found, at POD14, using stereology and a wide range of markers known to identify activated microglia and reactive astrocytes, that the maintenance of tumor-induced bone pain is dissociated from spinal microglia and astrocyte activation in MRMT-1-inoculated female rats. In contrast, several studies report the presence of microglial cells and astrocytes in the activation state within the spinal cord dorsal horn of cancer-bearing animals ^58–61^. These differences in spinal glial activation require further investigation, but species, strain and sex, type of cancer cells, site of tumor cell inoculation and the time window at which glial cells were studied may explain at least some of these discrepancies.

In summary, our results show that peripheral T cell infiltration into DRGs is of pivotal importance in cancer-induced bone pain and that selective allosteric inhibition of CCL2/CCR2 signaling by PP101pepducin directly and indirectly blocks the underlying inflammatory and neuropathic processes that lead to bone cancer pain.

## Methods

### Pepducin synthesis

Unless otherwise stated, all reactions were performed under nitrogen atmosphere. All solvents (ChemImpex International), coupling reagents (ChemImpex International) and N-methylpyrrolidinone (A&C American Chemicals Ltd) were HPLC grade. All other reagents were purchased from Sigma-Aldrich. Water sensitive reaction were performed in anhydrous solvents. All peptides were synthesized at a 0.1 mmol scale on an automated Symphony-X peptide synthesizer system (Gyros Protein Technologies) using 0.22 mmol/g Tentagel S RAM resin *S (Rapp Polymere*). Coupling of Fmoc amino acids or palmitic acid (ChemImpex International #35152) was performed in 1-[bis(dimethylamino)methylene]-1H-1,2,3-triazolo[4,5-b]pyridinium 3-oxide hexafluorophosphate (HATU, 5 fold excess) in the presence of *N*,*N*-diisopropylethylamine (DIPEA, 10 fold excess) twice for 30 min using conventional Fmoc-based solid-phase peptide synthesis (SPPS). The deprotection steps were performed using 20% piperidine (A&C American Chemicals Ltd) in dimethylformamide (DMF). Once all amino acid were coupled, the terminal Fmoc was removed and the peptides were cleaved from the polymer solid support using a mixture of trifluoroacetic acid (TFA)/water/triisopropylsilane(TIPS)/1,2-ethanedithiol (EDT) (92.5/2.5/2.5/2.5, v/v/v/v) under stirring for 3 h. The peptide mixtures were filtered, and the filtrates precipitated in methyl *tert*-butyl ether (MTBE). The precipitated crude peptides were centrifugated (3000 rpm, 10 min) and the ether layer removed by decantation. The crude peptides were dissolved in a mixture of water/acetonitrile (1:1, v/v) and lyophilised on a Freeze Zone lyophilizer (Labconco). Peptides were then dissolved in DMF, filtered, and purified with on a mass triggered preparative HPLC system (Waters) equipped with a X-select CSH C18 (5 µm, 10 x 100 mm) column. Pure fractions were lyophilized to give the final product as white powder. The peptides were further dissolved in water/acetonitrile 1:1 with 0.1 % TFA to obtain TFA salt. Peptides were analyzed using a Acquity UPLC-MS system class H (column Acquity UPLC protein BEH C4 (2.1 mm × 50 mm), 1.7 μm particles with pore 300 Å) coupled with a SQD2 mass spectrometer (Waters), and a PDA eλ UV-visible detector with the following gradient: 0 min, 5% acetonitrile (ACN); 0.2 min, 5% ACN; 1.5 min, 95% ACN; 1.8 min, 95% ACN; 2.0 min, 5% ACN; 2.5 min, 5% ACN. Flow: 800μL/min with 0.1% formic acid as buffer. The purity of all reported compounds was >97%.

For the synthesis of TAMRA-labeled pepducins, 2-aminohexadecanoic acid (Sigma-Aldrich #08051-1G) was protected as described by Koppitz et al. ^62^. The resulting Fmoc-2-aminohexadecanoic acid was used without further purification for elongation of the peptide. TAMRA carboxylic acid (5(6)-carboxytetramethylrhodamine, Sigma-Aldrich #C2734) was coupled at the N-terminus of the peptide using the same procedure as the Fmoc amino acid.

### Cell culture

MRMT-1 rat breast carcinoma cells were kindly provided by the *Cell Resource Center for Biomedical Research Institute of Development, Aging and Cancer* (Tohoku University) and were harvested in RPMI 1640 medium supplemented with 10% FBS and 2% penicillin/streptomycin. HEK293 and HEKCCR2 cells were harvested in DMEM medium supplemented with 10% FBS and 2% penicillin/streptomycin. HEK293 medium also contained 20 mM HEPES. The cells were maintained at 37°C and 5% CO_2_ in a humidified atmosphere.

### BRET-based cellular assay

BRET-based cellular assays were performed to monitor G-protein dissociation and β-arrestin recruitment in response to compound stimulation. In these cellular assays, 2 x 10^6^ HEK293 cells were seeded onto 100 mm^2^ cell culture dishes and transfected 24 hours later with a 12 μg cDNA mix prepared in Opti-MEM serum-free media with the polyethylenimine (PEI) (Polysciences #23966-2) transfection agent at a 3:1 ratio (PEI: DNA). For the BRET assays that monitored CCR2-mediated G-protein signaling, the cells were transfected with either of the following biosensor couples: rCCR2, Gα_i1_-RlucII, Gβ_2_, and GFP10-Gψ_1_; or rCCR2, Gα_oA_-RlucII, Gβ_1_, and GFP10-Gψ_1_. For the BRET assays that monitored G-protein signaling at other GPCRs, the rCCR2 construct was replaced by plasmids coding for NTS1, MOR or CXCR4. For the BRET assays monitoring β-arrestin 2 recruitment to CCR2, the cells were transfected with β-arrestin-2-RlucII (R393A) and rCCR2-GFP10 constructs. The BRET biosensors were kindly provided by Dr Michel Bouvier (Université de Montréal). Twenty-four hours post-transfection, cells were detached using trypsin-EDTA and were seeded into white opaque 96-well plates at a concentration of 50 000 cells/well. Forty-eight hours post-transfection, adhered cells were washed with PBS and stimulated with compounds prepared in a 100 μL volume of HBSS (20 mM HEPES). Two distinct protocols were used to measure either an *agonist* or an *antagonist* effect. In the first protocol (a time-course experiment), the cells were stimulated with 100 nM CCL2 (R&D Systems # 3144-JE-025/CF) or 10 mM pepducin, then stimulated with the coelenterazine 400A substrate (5 mM) (Gold Biotechnology #C-320-1) and read for 30 min on Mithras2 Plate Reader (Berthold) using a BRET2 filter set (400-450 nm and 500-550 nm emission filters). In the antagonist mode, cells were first stimulated with a range of pepducin concentrations (10 nM to 10 mM, half-log concentration intervals) and incubated for 10 min at 37°C. The cells then received an EC80 concentration of CCL2 (3 nM). For the cells transfected with other GPCRs (NTS1, MOR, CXCR4, CCR5), they instead received EC80 concentrations of neurotensin (NT(8-13)), DAMGO, SDF-1α or rCCL5, respectively. The cells were then stimulated with coelenterazine 400A (5 μM) and read on the Mithras2 Plate Reader 10 min later. Once the plates were read, BRET^2^ ratios were determined by dividing the GFP^10^-associated light emission (515 nm) by RlucII-colenterazine 400A-associated light emission (400 nm). In the time-course experiments, these ratios were subtracted from the baseline values to determine ΔBRET. In the endpoint experiments, the data was normalized relative to CCL2 (or to other native ligands); values for non-treated cells were set as 0% pathway activation, and those for cells treated with the EC80 CCL2 (or with the other native ligands) were set as 100% pathway activation. The data represent three distinct experiments, tested in triplicate.

### Animals

Female Sprague-Dawley rats (150-175 g) (Charles River Laboratories) were maintained on a 12 h light/dark cycle with access to food and water *ad libitum*. All animal procedures were approved by the ethical committee for animal care of the *Université de Sherbrooke*, in compliance with the policies and directives of the *Canadian Council on Animal Care* and with the ARRIVE guidelines.

### PP101 *in vivo* selectivity assay

Animals were randomly assigned to experimental group and injected intrathecally with saline, CCL2 (0.1 µg/rat), CCL2 + SCR101 or CCL2 + PP101 (125 nmol/rat) under light isoflurane anesthesia.

### Cancer implantation surgery

Rats were randomly assigned either to cancer or sham surgery groups. Syngeneic MRMT-1 breast cancer cells were surgically implanted as described by Doré-Savard *et al*. ^63^. Briefly, 30,000 cells were diluted in 20 µL Hank’s Balanced Salt Solution (HBSS) and injected into the medullary cavity of the female rat femur after a minimal opening by a microdrill. The hole was then sealed with dental amalgam. No post-operative analgesia was used to avoid interference with pain assessment. Sham-operated rats underwent all surgery procedures, but no cancer cells were delivered into femoral bone marrow.

### Pepducin administration

Eleven days after cancer cell implantation, animals were randomly assigned to control (SCR101) or treatment (PP101) group. Rats received a daily administration of either PP101 (125 nmol/rat in DMSO, intrathecally) or SCR101 starting day 11 until day 14 under light anesthesia. Injections were performed 45 min prior to behavioral testing. The dose of PP101 was selected based on a previous proof-of-concept *in vivo* pepducin study performed by our group ^64^. Based on their mode of action, where pepducins must passively flip into the inner leaflet of the cell membrane to reach their intracellular target, it is estimated that only a fraction of the compound interacts with the target receptor and therefore pepducins are known to be administered at much higher concentration than orthosteric drugs.

### Paw retroflexion score

Animal paw retroflexion (ipsilateral) was monitored according to the following criteria: 0 = full contact with floor, 1 = light contact with floor, 2 = paw deformation light contact with floor, 3 = curved paw with only tip toes touching the floor, 4 = curved paw no contact with the floor. See supplementary Fig.4 for representative photos. A higher score indicates more pain.

### Burrowing behavior assessment

Rats were acclimatized for three consecutive days to the experimental setup and trained to burrow. On the experimental day, rats were placed individually in a bedding-free cage containing a transparent plastic cylinder (32 cm length, 10 cm width, opening 6 cm in height maintained by two angled bolts) filled with 2000 g of soft gravel. Animals were let to burrow for two hours, then gravel burrowed out of the cylinder was weighted.

### Dynamic weight bearing

The dynamic weight bearing (DWB) device (Bioseb) consisted of a Plexiglas enclosure (22 × 22 ×30 cm) with a floor sensor composed of 44 × 44 captors (10.89 mm^2^ per captor). The animal was allowed to move freely within the apparatus for 5 min while the pressure data and live recording were transmitted to a laptop computer via a USB interface. Raw pressure and visual data were colligated with the DWB software v1.4.2.98.

### Mechanical allodynia

Rats were acclimatized for three consecutive days to the von Frey apparatus. The mid-plantar surface of the ipsilateral and contralateral hind paws was stimulated with von Frey hairs of logarithmically increasing stiffness by an experimenter blinded to treatment. The von Frey hairs were held for 7 sec with intervals of several seconds between each stimulation. Stimuli were presented in a consecutive fashion, descending when a behavioral response was observed, ascending otherwise. The test ended after a negative response on the stiffest hair (15 g), or four stimulations after the first positive response. The 50% paw withdrawal threshold was interpolated using the formula: 50% g threshold = 10^[Xf+kδ], where Xf = value (in log units) of the final von Frey hair used; k = tabular value for the pattern of positive/negative responses; and δ = mean difference (in log units) between stimuli.

### Ex vivo µCT

Anesthetized rats were intraaortically perfused with 100 ml saline followed by 500 ml 4% paraformaldehyde solution. Ipsilateral femurs were removed and post-fixated 48 h in the same solution then washed in PBS for *ex vivo* µCT experiment. Scans were performed at the *McGill Bone Center* using a high-resolution desktop Micro-CT scanner, as previously described ^65^. Rat femurs were scanned at X-ray source power of 45 keV/222 µA and at a resolution of 11.25 µm/pixel. The µCT images were reconstructed using NRecon (v1.6.1.3) and CT-Analyzer (v1.10.0.2) provided by SkyScan which was used for reconstruction and 3D analyses, respectively. The Volume of Interest (VOI) of cortical + trabecular bone is defined as the total (tissue) volume including cortical bones, trabecular bones and any spaces over the range of 5.626 mm (201 cross sections) starting from the growth plate in the distal femur. VOI for trabecular is defined as the total (tissue) volume including all trabecular bones and any spaces over the range of 5.626 mm (201 cross sections) starting from the growth plate in the distal femur.

### *Ex vivo* tumor measurement

Anesthetized rats were intra-aortically perfused with 100 ml saline followed by 500 ml 4% paraformaldehyde solution. Ipsilateral femurs were removed and post-fixated 48 h in the same solution then washed in PBS. Femurs were immerged in a water-filled plethysmometer (IITC Inc. Life Science #520MR and #520A) up to the femoral head. The bone volume from sham animals was subtracted from cancer animals to obtain the tumor volume.

### Immunostaining

After perfusion, L1 to L3 DRG and spinal cord sections were collected, post-fixed in 4% paraformaldehyde solution at 4°C for 24 h and then cryoprotected in 30% sucrose in PBS at 4°C for 48 h. Frozen tissues were embedded at −35°C in O.C.T. compound and 30 µm transverse spinal cord sections were generated using a Leica SM220R sliding microtome and 20 µm DRG sections were generated using a cryostat on gelatin-coated SuperFrost Plus slides. For spinal cord glial staining, sections were demasked in 0.01 M citrate buffer, 0.05% Tween-20 at pH 6, 80°C for 45 min. After 20 min incubation in 1% NaBH_4_ in PBS, sections were blocked (10% NGS, 1% BSA, 0.05% Tween-20, 0.1% Triton X-100 in PBS) and incubated in 0.3 M glycine containing 0.2% Tween 20. Sections were labeled with mouse anti-pERK (1:100, Cell Signaling #9101), mouse anti-CD45 (1:500, Bio-Rad #MCA43R), mouse anti-CD68 (1:500, Bio-Rad #MCA341R), mouse anti-CD11b (1:500, Millipore, #CBL1512), and mouse anti-NeuN (1:500, Millipore #MAB377), rabbit anti-CCL2 (1:100, PeproTech #500-P76), rabbit anti-connexin 43 (1:500, Sigma #6219), rabbit anti-GFAP (1:500, Sigma, #G3893), rabbit anti-Iba1 (1:500, Wako #019–19741), rabbit anti-pP38 (1:100, Cell Signaling, #4511), rabbit anti-Ki67 (1:100, Abcam #AB16667), and mouse anti-pJNK (1:100, Cell Signaling #9251), chicken anti-CCR2 (1:200, Aves Labs, custom) and chicken anti-CGRP (1:500, Millipore #AB5705), guinea-pig anti-substance P (1:500, Neuromics #GP14110), hamster anti-CD3 (1:500, Santa Cruz, #sc-1174) in blocking buffer for three days at 37°C for spinal cord, 24 hour at R.T. for DRG. They were further incubated with fluorophore-conjugated secondary antibodies (1:500, AlexaFluor 488, 568, 647, Invitrogen) and spinal cord sections were mounted on SuperFrost Plus slides. All slides were coverslipped with ProLong Diamond Mountant. pERK^+^, CCL2^+^ and CCR2^+^ neurons and Iba1^+^ cells, were manually counted using ImageJ software, as well as fluorescence intensity quantifications. Neurons were considered CCL2^+^ when both membrane and cytoplasmic vesicle staining were present. Iba1^+^CD68^+^ and CD3^+^/DAPI^+^ cells were manually counted using Neurolucida 360 software. T cells were counted only if the CD3^+^ staining surrounded a large DAPI^+^ nuclei. For immunohistochemistry, sections were incubated in 0.3% H_2_O_2_ for 1 hour and blocked (3% NGS, 0.3% Triton X-100 in PBS). Sections were incubated in rabbit anti-C-Fos (1:5000, Abcam, #AB7963) in 1% NGS, 0.3% Triton X-100 (24 hours, 37°C) followed by peroxidase-conjugated secondary antibody (1:200 biotinylated anti-rabbit Vector Labs #BA-1000) and incubated in Elite ABC solution. The product of immune reaction was revealed using 3,3′-diaminobenzidine (DAB) as a chromogen and 0.015% H_2_O_2_. Finally, slices were dehydrated in graded ethanol, defatted in xylene and mounted with Permount. Fluorescence images of spinal cord and DRG slices were acquired at 20x using a Leica DM4000 microscope equipped with a Leica DFC350FX or a Zeiss AxioImager M2 equipped with a Hamamatsu Flash camera using the same acquisition parameters. For c-Fos immunoreactive nuclei quantification, positive neurons were counted after acquisition using a bright-field microscope and an InfinityX camera. C-fos-positive neurons were counted in spinal lamina I on the ipsilateral side which was delineated according to Molander. Neurons were considered positive if the nucleus showed the characteristic staining of oxidized DAB and was distinct from the background. Quantification was performed using ImageJ software. 5-10 sections located between vertebrae L1-L3 were analyzed per animal, for a total of three animals per treatment condition. Representative fluorescence images were acquired using an Olympus FV1000 confocal microscope and whole slide images were acquired using Zeiss AxioImager M2 equipped with a Hamamatsu Flash camera. All antibodies were validated on positive tissues and specificity controls were performed by omitting primary or secondary antibodies, or both. See supplementary Fig. 2 for representative photos of antibody specificity staining validation.

### PP101 penetration into DRG

PP101-TAMRA (125 nmol/rat), or vehicle (DMSO) were injected intrathecally in naïve animals. 45 min post-injection, anesthetized rats were decapitated and DRG of L6 to T12 lumbar vertebrae were isolated, post-fixed overnight in 4% paraformaldehyde at 4℃, and then cryoprotected in a 30% sucrose in PBS at 4℃. Frozen tissues were embedded in O.C.T. compound and 15 µm DRG sections were generated using a cryostat. Sections were washed twice in PBS and incubated with DAPI staining (1:8000 dilution) overnight at RT. They were then rinsed twice with PBS for 10 min, mounted on SuperFrost Plus slides and coverslipped with Prolong Diamond Antifade Mountant. Images were acquired on an Olympus FV1000 confocal microscope.

### Isolation of DRG macrophages and T cells

Anesthetized rats were decapitated and L1-L3 DRG tissue were rapidly dissected. DRG tissues were digested for 50 min with 0.025 % collagenase A (cat no. 10103586001 Roche), 1 mM EDTA in PBS at 37°C under constant agitation and triturated at 20, 40 and 50 min. Cells were then filtered through a 70 µm cell strainer (Fisherbrand #22363548) and washed with FACS buffer (10% NGS, previously decomplemented at 56°C, 30 min + 1 mM EDTA in PBS). Cells were centrifuged at 1600 RPM at 4°C and resuspended in FACS buffer.

### Flow cytometric assays

Dissociated cells were briefly fixed with 2% paraformaldehyde for 30 min at 4°C, centrifuged at 1600 RPM at 4°C, then washed with cold FACS staining buffer. Cells were centrifuged at 1600 RPM at 4°C, resuspended in 100 μl of FACS buffer and incubated with anti-CD32 antibody (1:20, BD Biosciences #550271) for 30 min at 4°C to block nonspecific Fc receptor binding. Multicolor immunostaining was performed with F4/80-AlexaFluor 647 (1:40, Bioss Antibodies #BS-7058R-A647), CD163-FITC (1:40, Biorbyt #orb434303), CD68-PE-Cy7 (1:40, Abcore #AC12-0231-17), CD3-APC-Cy5.5 (1:40, Abcore #AC12-0212-04), CD4-PerCP-eFluor710 (1:40, eBioscience #46-0040-82) and CD8a-PE (1:40, BioLegend #200607) antibodies in FACS buffer with 2 µg DAPI to distinguish cells from debris, for 1h at 4°C. Cells were rinsed, resuspended in FACS buffer containing DAPI and analyzed using a Beckman Coulter CytoFLEX flow cytometer (Beckman Coulter, Brea, CA, USA). The specificity of each antibody was validated on activated rat splenocytes and peritoneal macrophages (intraperitoneal thioglycollate 2%, 3mL, 72 h) isolated as described in the supplementary material section. Single-stained compensation beads (VersaComp Antibody Capture Beads #B22804) were used to measure the single dye fluorescence spill over to neighborhood channels for multicolor compensation. An electronic compensation matrix was used to correct this crosstalk between channels. Positive and negative selection gates were set using fluorescence minus unstained cells. Fluorescence intensity distribution was analyzed with CytExpert software (Beckman Coulter).

### T cell depletion strategy

Cancer-implanted animals were injected i.t. with anti-CD4 (2 µg/rat dissolved in saline solution, clone OX-38, InVivoMab #BE0308), anti-CD8 (2 µg/rat dissolved in saline solution, clone OX-8, Bio-Rad #MCA48G) or anti-rat TCRαβ antibody (2 µg/rat dissolved in saline solution, clone R73, Bio-Rad #MCA453GA) from day 9 to day 14 to deplete DRG T cells as previously described ^66^. An equal dose of isotype rat IgG1 (Bio-Rad #MCA1209) or rat IgG2a (Bio-Rad #MCA1210) antibody was administered as control.

### Glial cell morphology analysis

Iba1+ microglia and GFAP+ astrocytes were acquired using a Zeiss AxioImager M2 microscope attached to a Hamamatsu Flash 4.0 V2+ CMOS camera, an ApoTome 2 system and a three-axis motorized stage. Cells were analyzed in systematic random sampling (SRS) using StereoInvestigator 2018.1.1 software. The regions of interest (ROI) (ipsilateral lamina 1-3) were outlined using a 5× objective according to the cytoarchitectonic organization of the spinal cord; cells were counted using a 63× oil-objective lens. The sampling grid was randomly distributed on the slice, such that the counting frames were placed randomly onto ROI. Cells were excluded if they were located on the edge of the tissue or outside of the counting frame. Microglial cell soma and processes were estimated using the nucleator method and the total number of microglia and astrocytes was estimated using the optical fractionator method. For each method, a counting frame size of 100 µm × 100 µm and a grid size of 200 µm × 200 µm were used, with a guard zone of 2 µm and a dissector height of 14 µm. Over 177 cells were analyzed per condition over three animals such as the error coefficient was below 0.15 for all animals.

### BSCB permeability assessment

Sodium fluorescein (NaFlu) was solubilized in sterile saline and injected intravenously (10%, 2 mL/kg), as previously described ^41^. After 30 min, rats were intra-aortically perfused with 100 mL saline and both DRG and spinal cord were collected and weighted. Samples were homogenized in PBS, followed by addition of an equal volume of 60% trichloroacetic acid to precipitate proteins. Samples were vortexed for 2 min, incubated for 30 min on ice, then centrifuged at 14,000 × *g* for 10 min at 4°C. Concentration of NaFlu in supernatant was quantified using a GENios Pro plate reader (440 nm ex. and 525-550 nm em.) and calculated using a standard curve.

### Statistical analysis

Statistical analysis was performed using GraphPad Prism 8.1.2 software. Data are presented as mean ± SEM. Differences between two groups were assessed using a Mann-Whitney test. Statistical analysis for multiple comparisons were performed using Kruskal-Wallis test followed by Dunn’s multiple comparison test or a Two-Way ANOVA followed by Sidak’s multiple comparison test. Hashtags represent differences with the sham + SCR101 group, *p* < 0.05 (#), *p* < 0.01 (##), *p* < 0.001 (###) *p* < 0.05 and stars represent differences with the cancer + SCR101 group, *p* < 0.05 (*), *p* < 0.01 (**), *p* < 0.001 (***).

## Author contributions

E.M. designed the project and carried out all experiments and analyzed results, with the exception of BRET-based assays. R.B. performed BRET-based experiments and analyzed results. C.M. and V.Z. synthesized pepducins. E.M wrote the manuscript with critical revisions provided by P.S., J.C. and S.W.K. P.S. acquired funding. All authors approved the final version of the manuscript.

## Acknowledgement

The BRET-based biosensors were generously provided by Michel Bouvier (Department of Biochemistry and IRIC, Université de Montréal, Montréal, QC, Canada), as a member of the CQDM team (M. Bouvier, T. Hébert, S.A. Laporte, G. Pineyro, J.-C. Tardif, E. Thorin and R. Leduc). We wish to thank Leonid Volkov for immune cell flow cytometry guidance and Michael Desgagné for improvement of pepducin solubility. We further wish to thank Allan Basbaum for critical revisions.

## Supplementary Figures

**Supplementary Figure 1:**
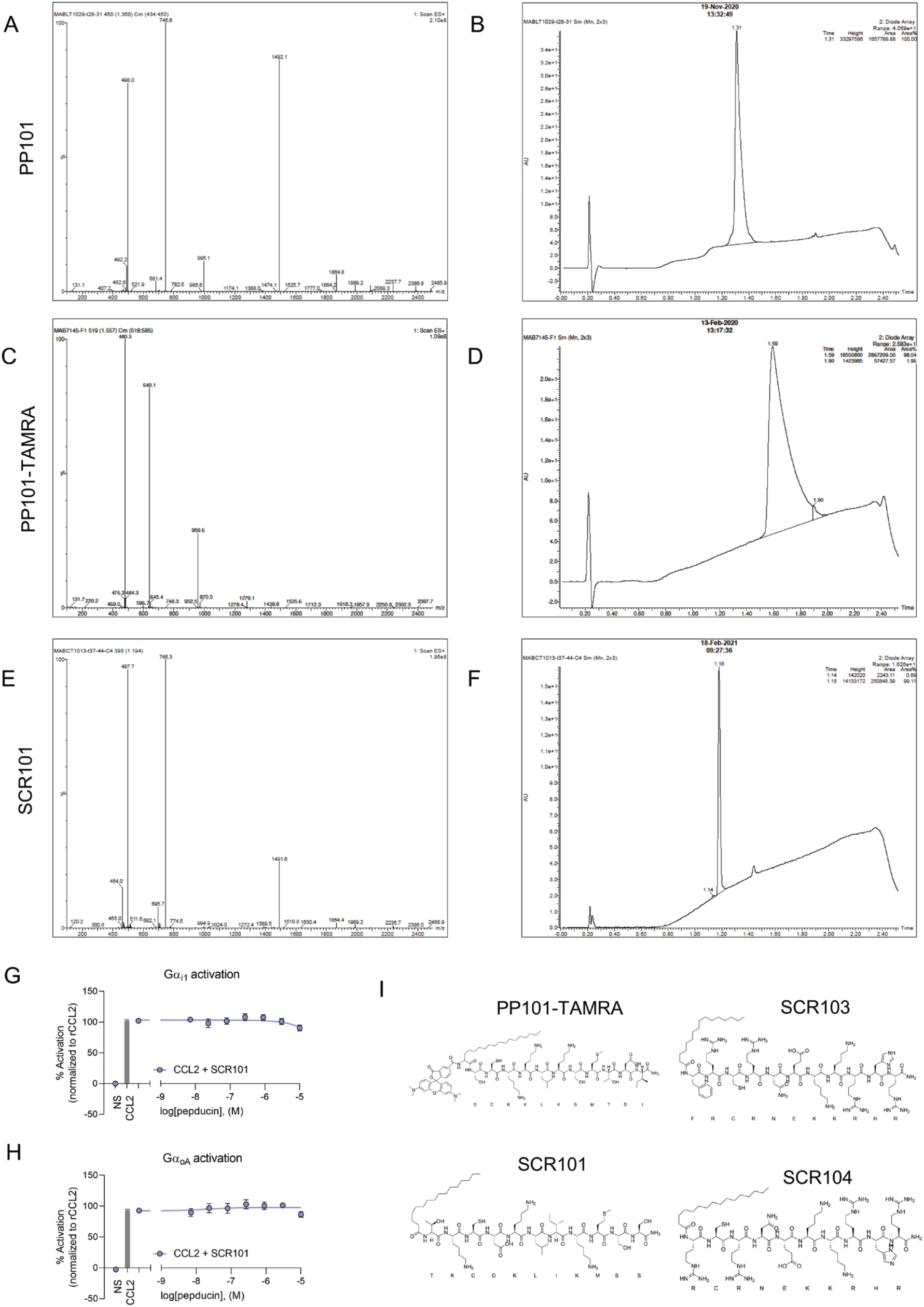
Effect of scrambled control CCR2 pepducins on CCL2-induced G-protein activation. (**A, C and E**) Mass spectrometry and (**B, D and F**) Ultra Performance Liquid Chromatography (UPLC) spectra of PP101, PP101-TAMRA and SCR101 peptides. Effect of the scrambled control pepducin, SCR101 on (**G**) Gαi1- and (**H**) GαoA-protein activation induced by 2 µM CCL2 in HEK293 cells transiently transfected with rCCR2 and BRET^2^-based biosensors (*n* = 3 duplicate). BRET^2^ ratios were normalized according to CCL2; values for non-treated cells were set as 0% activation, and those for cells stimulated with 2 µM CCL2 were set as 100% activation. (**I**) Amino acid sequences of the fluorescently labeled PP101-TAMRA pepducin and of the scrambled controls, SCR101, SCR103, and SCR104 used in the present study. Data represent mean ± SEM.

**Supplementary Figure 2:**
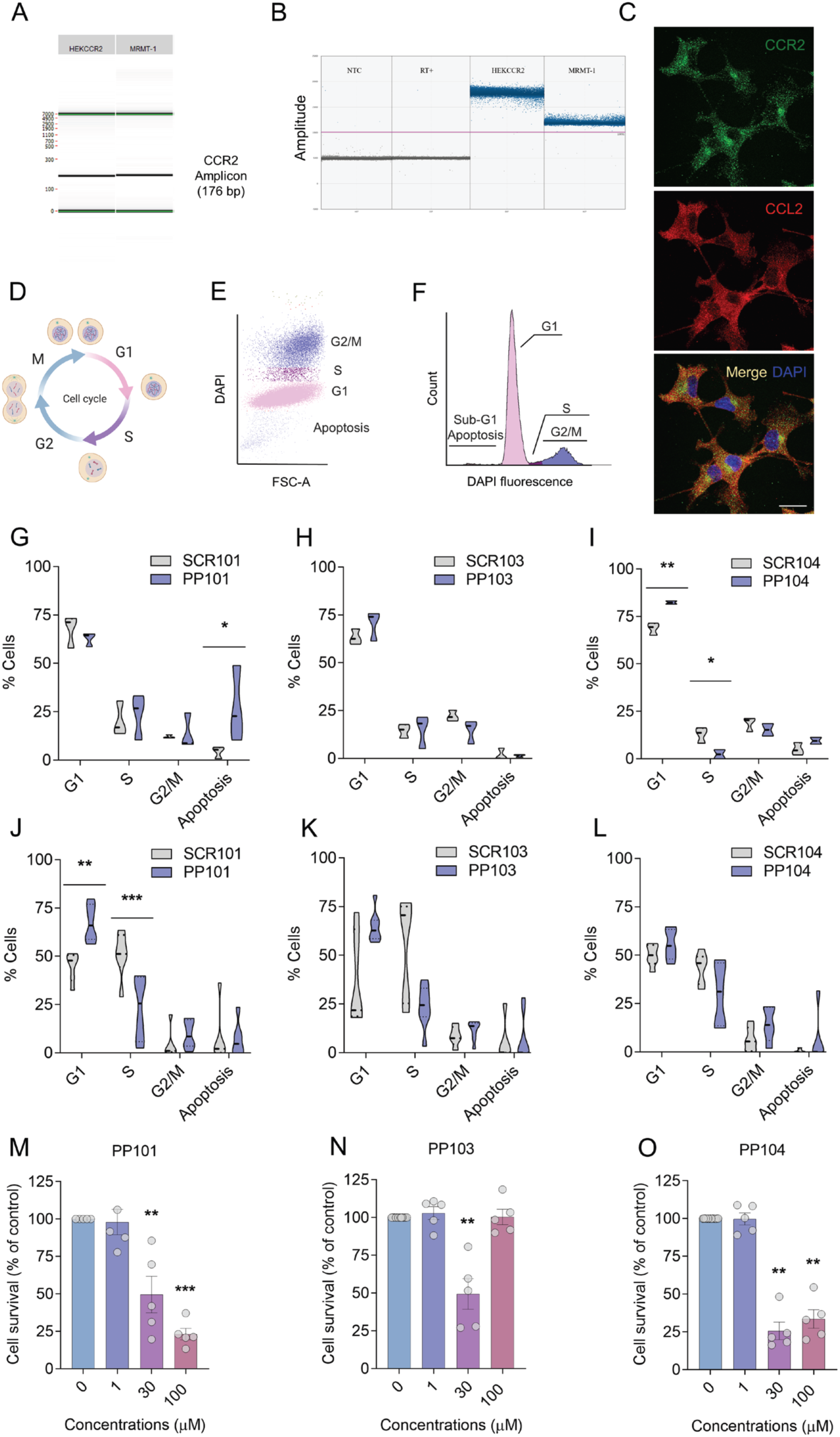
Pepducin antagonist efficacy *in vitro* on CCR2-expressing cancer cells. (**A**) RT-qPCR followed by capillary electrophoresis and (**B**) digital droplet PCR (ddPCR) analyses of CCR2 receptor mRNA expression in MRMT-1 breast cancer cells. RT-qPCR (*n* = 3) and ddPCR (*n*=4). In (**B**), each gray dot represents a single negative amplification reaction of CCR2 complementary DNA (cDNA) and each blue dot represents a single positive amplification reaction of CCR2 cDNA. NTC: No coding template, controls the qPCR reaction; no cDNA with primers, RT+: Reverse transcriptase +, controls the reverse transcriptase reaction; no mRNA with RT enzyme. (**C**) Immunofluorescence analysis of both CCR2 and CCL2 protein expression in MRMT-1 cells. Scale bar corresponds to 20 µm. (**D-F**) MRMT-1 cell cycle was analyzed by flow cytometry using DAPI as the fluorescence DNA dye. FSC-A: Forward cell scatter area. Effect of PP101, PP103, and PP104 (100 µM) on MRMT-1 cell cycle after 12h (**G-I**; *n* = 3) and 20h post-incubation (**J-L**; *n* = 4-6), as compared to their respective scrambled control pepducins, SCR101, SCR103, and SCR104. (**M-O**) MRMT-1 cell viability assessed after 72h incubation with PP101, PP103 and PP104 using the WST-1 assay (*n* = 5 triplicate). Data represented as mean ± SEM. Two-way ANOVA followed by Sidak’s multiple comparisons test in (**G-L**), Kruskal-Wallis followed by Dunn’s multiple comparisons test in (**M-O**) and Mann-Whitney test in (**P**). \**P* < 0.05, \*\**P* < 0.01, \*\*\**P* < 0.001.

**Supplementary Figure 3:**
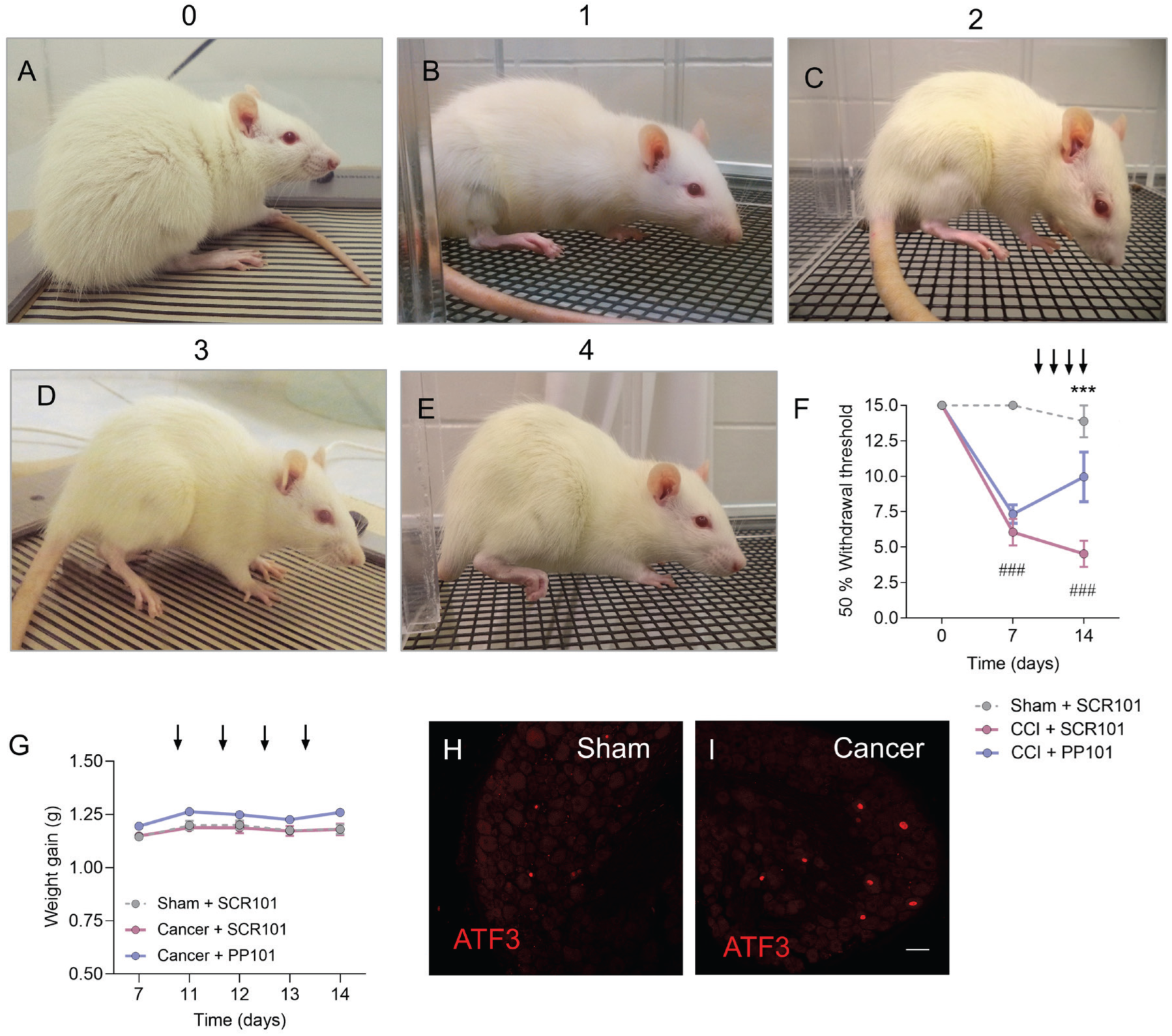
Pain-related behaviors associated to neuropathic and bone cancer pain. Changes in paw retroflexion scores, a coping behavior used to prevent the pain generated by the injured tumor-bearing paw pressure against the floor, were recorded as follows (**A**) score 0 - full contact with floor, (**B**) score 1 - light contact with floor, (**C**) score 2 - paw deformation with light contact with floor, (**D**) score 3 - curved paw with only tip toes touching the ground, (**E**) score 4 - curved paw with no contact with the ground. The higher the retroflexion score, the more pain. (**F**) Repeated daily administrations of PP101 or SCR101 (125 nmol/rat, i.t.) between days 11 and 14 following chronic constriction injury (CCI) (sham+SCR101 *n* = 5, CCI+SCR101 *n* = 8, CCI+PP101 *n* = 6). (**G**) Repeated daily administrations of PP101 or SCR101 (125 nmol/rat, i.t.) between days 11 and 14 on animal weight gain (sham+SCR101 *n* = 7, cancer+SCR101 *n* = 7, cancer+PP101 *n* = 6). (**H**, **I**) Immunostaining of the neuronal injury marker ATF3 in DRG neurons of sham and tumor-bearing rats.

**Supplementary Figure 4:**
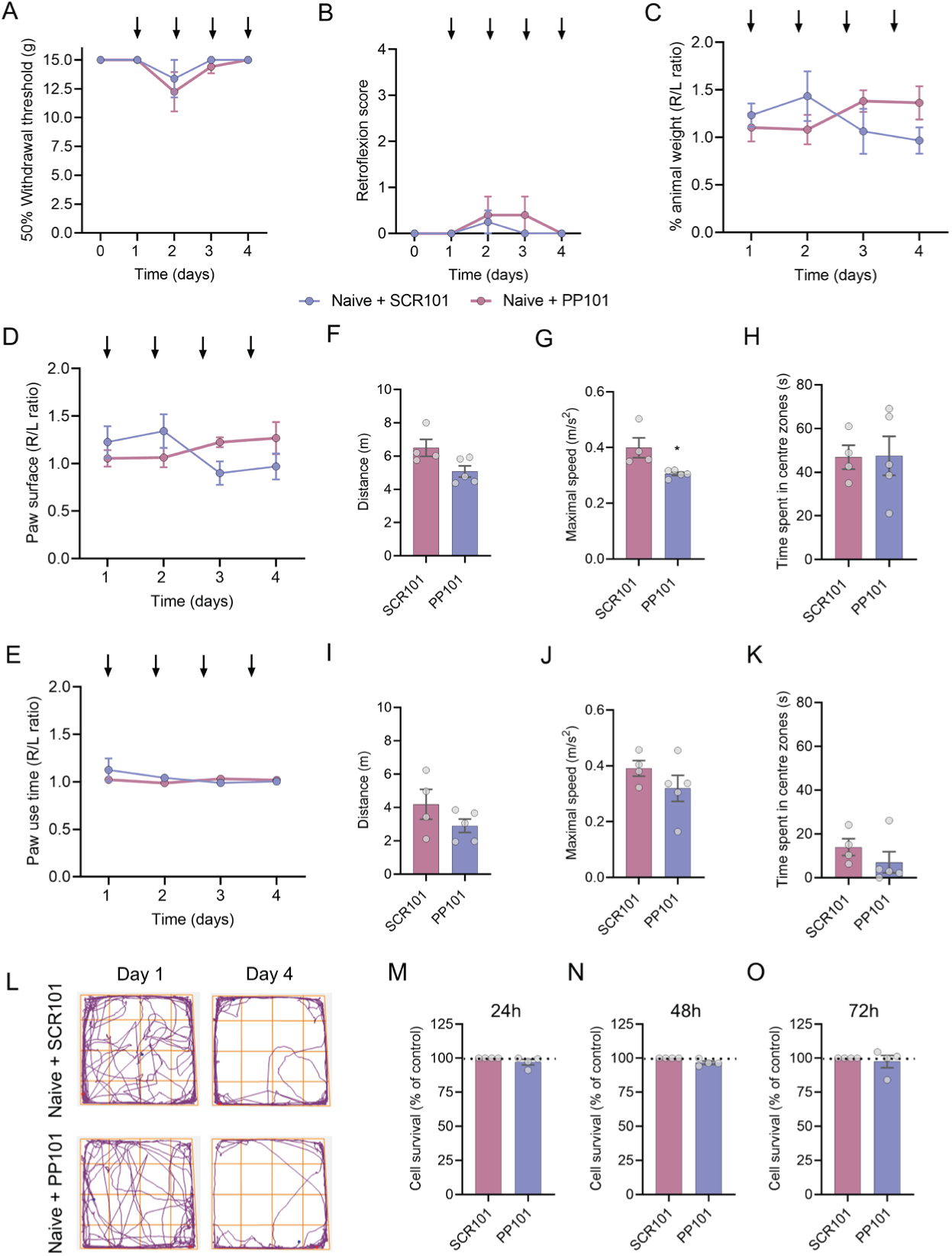
Lack of sensory, motor and anxiety-like adverse effects of PP101 or SCR101 in naïve rats and absence of toxicity on DRG neurons. Repeated daily administrations of PP101 or SCR101 (125 nmol/rat, i.t.) between days 1 and 4 on (**A**) mechanical thresholds (**B**) paw retroflexion score and (**C**, **D** and **E**) dynamic weight bearing in naïve animals. (**F**) Distance, (**G**) maximal speed and (**H**) anxiety-like behaviors in the open field test following a single administration of PP101 or SCR101 (125 nmol/rat, i.t.). (**I-L**) Distance, maximal speed, and anxiety-like behaviors in the open field test following four daily administrations of PP101 or SCR101 (125 nmol/rat, i.t.) (SCR101 *n* = 4, PP101 *n* = 5). (**M-O**) Cell viability of the immortalized DRG cell line F11 assessed after 24h, 48h and 72h incubation with PP101, using the WST-1 assay (*n* = 1 quadruplicate).

**Supplementary Figure 5:**
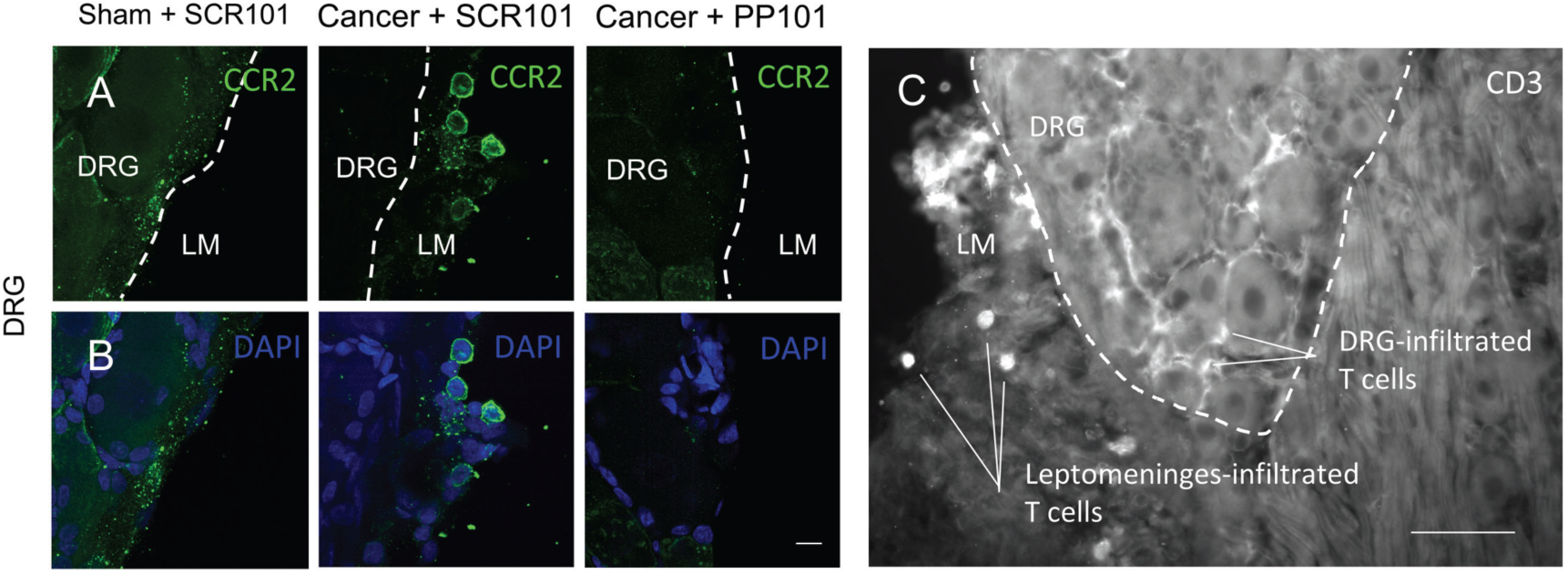
Infiltrating CCR2^+^ immune cells into the DRG leptomeninges and CD3^+^ T cells into the DRG parenchyma. (**A-B**) CCR2^+^ immune cells in the DRG leptomeninges of cancer-bearing animals. Dash lines delineating DRG and surrounding leptomeninges. DRG: Dorsal root ganglia, LM: Leptomeninges. All scale bars correspond to 10 µm. (**C**) CD3^+^ T cells in the DRG leptomeninges and parenchyma of cancer-bearing animals. Scale bar corresponds to 150 µm. Scale bar corresponds to 100 µm.

**Supplementary Figure 6:**
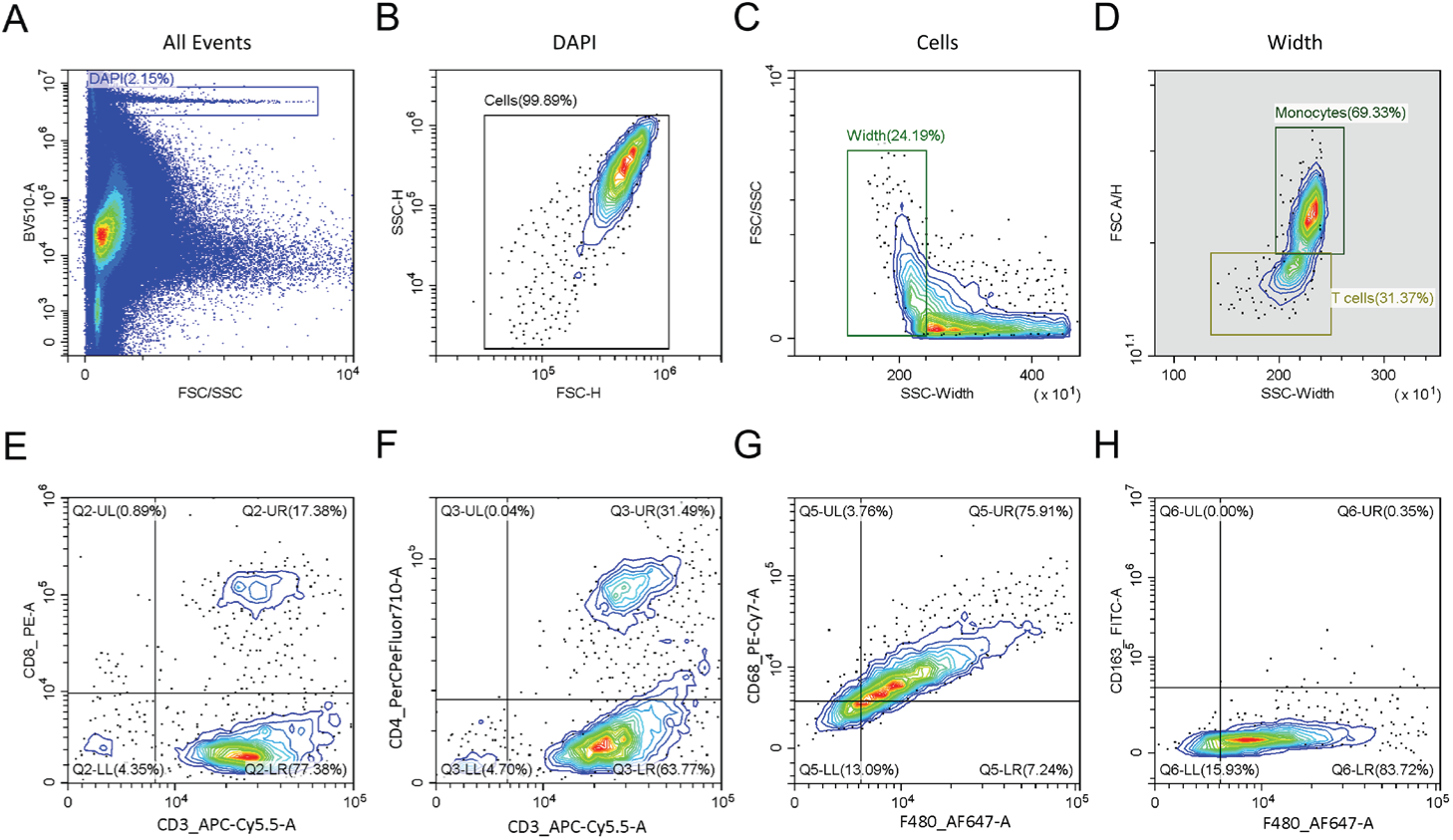
(**A-D**) The gating strategy used for the analysis of CD4^+^ and CD8^+^ T cell and CD68^+^ and CD163^+^ DRG infiltrating monocytes are illustrated by the gates. Splenocytes and peritoneal macrophages were isolated from a thioglycollate-induced peritonitis model in rats, pooled and used as positive control for activated immune cells. The cell suspension was stained with all antibodies. Doublets and debris were discriminated on a BV510-A and FSC/SSC plot. Cell width was gated on a FSC/SSC and SSC-Width plot. (**E and F**) T cells and monocyte populations were gated on a FSC A/H and SSC-Width plot. The CD8^+^ and CD4^+^ T cells were identified on CD8 and CD3 and CD4 and CD3 plots, respectively. (**G and H**) The CD68^+^ and CD163^+^ monocytes were identified on a CD68 and F4/80, and CD163 and F4/80 plots, respectively. BV: bright violet, FSC: forward scatter, SSC: side scatter, H: height, A: area, CD: cluster of differentiation, PE: phycoerythrin, AF: AlexaFluor, PerCP: peridinin chlorophyll, Cy-: cyanine, PerCP: peridinin-chlorophyll-protein complex, FITC: fluorescein isothiocyanate.

**Supplementary Figure 7:**
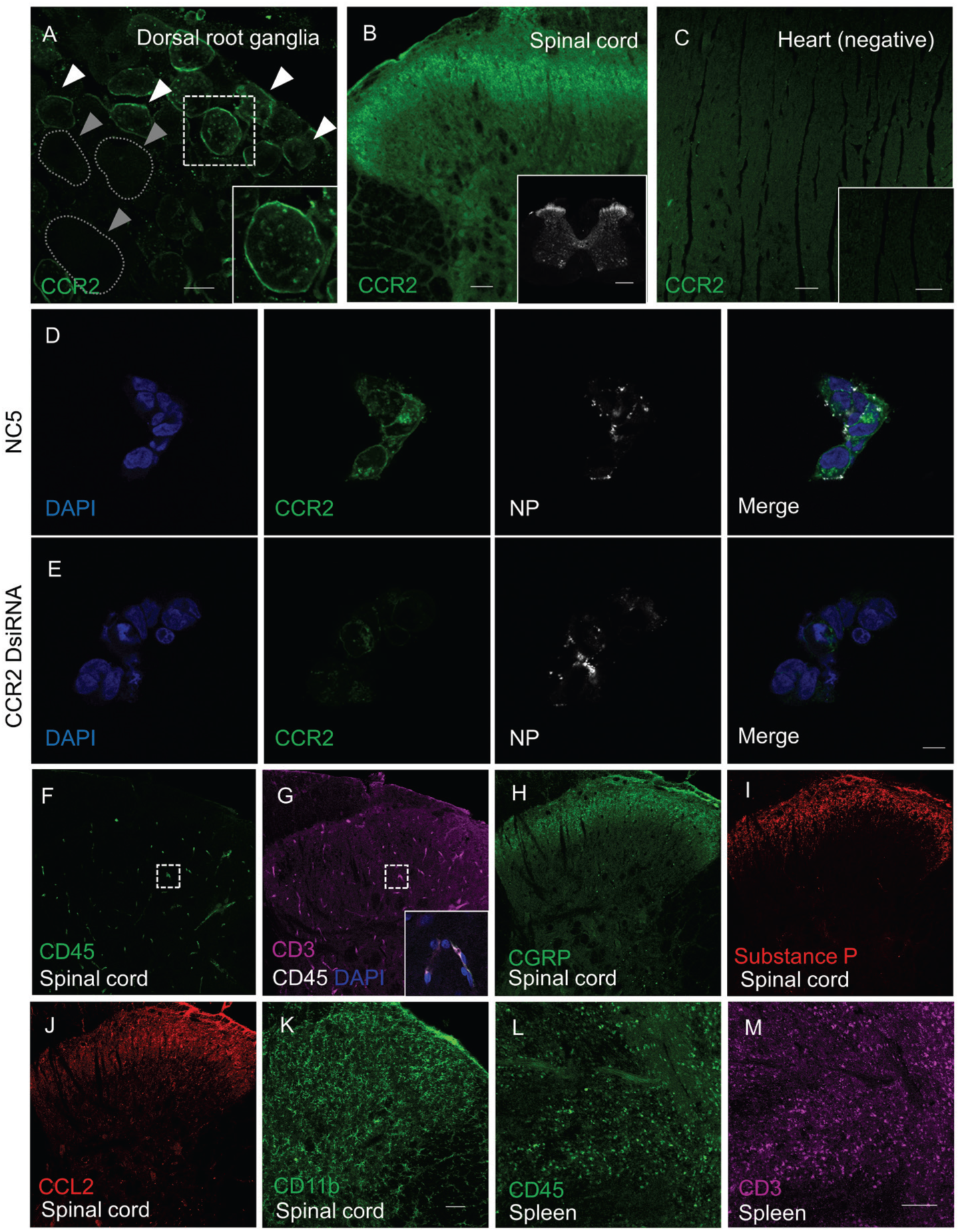
Antibody specificity. A custom-made anti-rat CCR2 antibody (Aves Labs) was produced, and its specificity validated in (**A**) DRG and (**B**) spinal cord used here as CCR2-positive tissue as well as in heart serving as negative control tissue (**C**). White arrows represent CCR2 staining in small and medium neurons and gray arrows represent CCR2-negative neurons. Scale bars correspond to 20 µm in (**A**), 100 and 500 µm (framed image) in (**B**) and 50 µm in (**C**). (**D**, **E**) HEK cells expressing the CCR2 receptor were treated with Neuro9 lipid nanoparticles encapsulating CCR2-targeting Dicer substrate siRNA or NC5, a non-targeting control siRNA ^67^. Scale bar corresponds to 10 µm. NP: DiD-tagged nanoparticles. Validation of the staining of CD45^+^ immune cells and CD3^+^ T cells in the spinal cord (**F-G**) of naïve animals, 24 hours following the second intrathecal injection of freshly isolated peritoneal macrophages and splenocytes. Magnified image of the spinal cord shows colocalization of CD45 and CD3. These immune cells seem to be colocalized within the spinal cord blood vessels rather than in the parenchyma. Validation of the staining specificity of (**H**) CGRP, (**I**) substance P, (**J**) CCL2 and (**K**) CD11b on spinal cord sections and (**L-M**) CD45 and CD3 in spleen sections of naïve animals. Scale bars correspond to 100 µm.

## Supplementary Material and Methods

### Quantitative RT-PCR (Supplementary Fig. 2)

MRMT-1 cell mRNA was extracted using RNeasy Mini Kit following the manufacturer’s protocol. RNA integrity was assessed using an Agilent 2100 Bioanalyzer. RNA quality and presence of contaminating genomic DNA were also verified. Reverse transcription was performed using 1 µg total RNA with QuantiTect reverse transcriptase, random hexamers, and dNTPs in a total volume of 20 µl. Forward and reverse primers were individually resuspended in Tris-EDTA buffer and diluted as a primer pair to 1 μM in RNase DNase-free water. Primer sequences for CCR2 are as follow: forward TGATCAGCACACTTGTGGCCC, reverse GCCTCACAGCCCTATGCCTC. qPCR reactions were performed in a volume of 10 µl in 96-well plates on a CFX96 Thermal Cycler with 5 μL of 2X iTaq Universal SYBR Green Supermix, 10 ng cDNA and 200 nM primer pair solution. The following cycling conditions were used: 3 min at 95°C followed by 50 cycles of 15 s at 95°C, 30 s at 60°C, 30 s at 72°C. The amplified products were analyzed by automated chip-based microcapillary electrophoresis on a Caliper LabChip 90 instrument. Amplicon sizing and relative quantitation was performed by the manufacturer’s software, before being uploaded to the LIMS database.

### Digital droplet PCR (Supplementary Fig. 2)

Droplet Digital PCR (ddPCR) reactions are composed of 10 µl of 2X QX200 ddPCR EvaGreen Supermix (Bio-Rad), 10 ng cDNA, 100 nM primer pair solution and molecular grade sterile water (Wisent) for a 20 µl total reaction. Each reaction mix (20 µl) was converted to droplets with the QX200 droplet generator. Droplet-partitioned samples were then transferred to a 96-well plate, sealed and cycled in a C1000 deep well Thermocycler under the following cycling protocol: 95°C for 5 min, followed by 50 cycles of 30 s at 95°C, 1 min at 59°C and 30 s at 72°C followed by post-cycling steps of 5 min at 4°C and 5 min at 90°C and an infinite 12°C hold. The cycled plate was then transferred and read using the QX200 reader. The concentration reported is in copies/µl of the final 1x ddPCR reaction (using QuantaSoft software from Bio-Rad).

### Analysis of cell cycle by flow cytometry (Supplementary Fig. 2)

30,000 MRMT-1 cells were seeded in triplicate in a 48-well plate in RPMI 1640 medium supplemented with 10% FBS and 2% penicillin/streptomycin. 12 h after cell seeding, cells were starved in serum-deprived medium during 24h for cell synchronization. Fresh medium containing 2% FBS with 100 µM of pepducin (0.4% DMSO), or their corresponding scrambled control was added. After 12 or 20h drug incubation, cells were detached using Accutase and inhibited with medium containing 2% FBS. Cells were pelleted using a microcentrifuge and the pellet was resuspended in 50 µL of FACS buffer (PBS containing 2% FBS). Cells were fixed with 70% ethanol at -20°C added in a dropwise manner while gently vortexing and then incubated on ice for 30 min. Cells were pelleted and resuspended in 150 µL FACS buffer containing DAPI at a concentration of 2 µg/mL and incubated on ice for 30 min. DAPI fluorescence was analyzed using a Beckman Coulter’s Cytoflex 30 flow cytometer at a flow rate of 30 µL/min in blue channel. Percentage of cells in the different phases of the cell cycle was calculated using ModFit software.

### WST-1 assay (Supplementary Fig. 2)

15,000 MRMT-1 cells were seeded in triplicate in a 96-well plate in RPMI 1640 medium without phenol red supplemented with 10% FBS and 2% penicillin/streptomycin. 12 h after cell seeding, 100 µl of fresh serum- and phenol red-free medium containing 100 µM of pepducin (0.4% DMSO), or corresponding scrambled control was added. After 72 h drug incubation, 10 µl of WST-1 reagent (5 mg/ml) was added to each well and incubated for 4h at 37 °C, 5% CO_2_. The optical density (OD) of the formazan dye generated by the conversion of the WST-1 tetrazolium salt by mitochondrial enzymes was measured at 450 nm for each well using a GENios Pro plate reader (Tecan). Cell viability was measured using the following formula: (OD_450nm_ treatment/OD_450nm_ scrambled control) × 100.

### CCR2 knockdown in HEK cells expressing CCR2 (Supplementary Fig. 6)

Cells were plated on glass coverslips (Fisherbrand #12-545-80) in DMEM medium supplemented with 10% FBS, 20 nM HEPES and 2% penicillin/streptomycin. 12 hours later, cells were treated with 1 mg/mL recombinant ApoE in addition to 200 nM scrambled control oligonucleotide or CCR2-targeting DsiRNA (IDT, IA, USA) encapsulated in Neuro9 lipid nanoparticles using a NanoAssemblr (Precision Nanosystems #NIS0003, BC, Canada). After 48 hours, cells were fixed with 4% paraformaldehyde solution and stained for CCR2 receptor using our custom-made chicken anti-rat CCR2 antibody (1:200, Aves Labs, CA, USA) and further incubated with fluorophore-conjugated secondary antibodies (1:500, AlexaFluor 488, Invitrogen), then mounted on SuperFrost Plus slides and coverslipped with ProLong Diamond Mountant (Invitrogen # P36970). Fluorescence images were acquired using an Olympus FV1000 confocal microscope.

### Intrathecal injection of peritoneal macrophages and splenocytes (Supplementary Fig. 6)

Anesthetized rats were euthanized, and peritoneal macrophages were collected by washing the abdominal cavity with PBS and collecting the cell suspension. Cells were centrifuged 5 minutes at 1600 RPM at 4°C and the cell pellet was resuspended in erythrocyte lysis buffer (RBC buffer) for 4 minutes at 4°C. Erythrocyte lysis was stopped by adding PBS. Cells were centrifuged and resuspended in PBS and filtered through a 70 µM cell strainer (Fisherbrand #22363548). Splenocytes were obtained by crushing a spleen through a 70 µM cell strainer using a sterile pestle and collecting the cells using PBS. Cells were centrifuged for 5 minutes at 1600 RPM at 4°C and the pellet was resuspended in RBC buffer for 4 minutes at 4°C. Erythrocyte lysis was stopped by adding PBS. Cells were centrifuged and resuspended in PBS and filtered through a 70 µM cell strainer. Macrophages and splenocytes were pooled and injected intrathecally in a naïve rat under light anesthesia. A second injection of freshly isolated macrophages and splenocytes was performed after 24 hours. 24 hours later, anesthetized rat was intraaortically perfused with 100 mL saline followed by 500 mL of freshly prepared 4% paraformaldehyde solution. L1 to L3 spinal cord sections and the spleen were collected, post-fixed in 4% paraformaldehyde solution at 4 °C for 24 h and then cryoprotected in 30% sucrose solution in PBS at 4 °C for 48 h. Frozen tissue were embedded at −35 °C in O.C.T. compound and 30 µm transverse spinal cord sections were generated using a Leica SM220R sliding microtome and 20 µm DRG sections were generated using a cryostat (Leica CM1860UV, Leica Biosystems, Concord, ON, Canada). Sections were labeled with mouse anti-rat CD45 (1:500, Bio-Rad #MCA43R) and hamster anti-CD3 (1:500, Santa Cruz, #sc-1174), and were further incubated with fluorophore-conjugated secondary antibodies (1:500, AlexaFluor 488, 647 Invitrogen) and mounted on SuperFrost Plus slides and coverslipped with ProLong Diamond Mountant (Invitrogen # P36970). Fluorescence images were acquired using an Olympus FV1000 confocal microscope.

### Chronic constriction injury model (Supplementary Fig. 2)

Female rats were randomly assigned either to CCI or sham surgery groups. Chronic constriction injury (CCI) was performed as previously described by Bennett and Xie ^68^ with a modification in the suture used (5-0 Prolene; Ethicon, Somerville, NJ, USA) to minimize suture-induced inflammation.

### ATF3 immunostaining (Supplementary Fig. 2)

Anesthetized rats were intraaortically perfused with 100 mL saline followed by 500 mL of freshly prepared 4% paraformaldehyde solution. L1 to L3 spinal cord sections and the spleen were collected, post-fixed in 4% paraformaldehyde solution at 4°C for 24 h and then cryoprotected in 30% sucrose solution in PBS at 4°C for 48 h. Frozen tissues were embedded at −35°C in O.C.T. compound and 15 µm DRG sections were generated using a cryostat (Leica CM1860UV, Leica Biosystems, Concord, ON, Canada). Sections were labeled with rabbit anti-rat ATF3 (1:200, HPA001562, Sigma) and were further incubated with fluorophore-conjugated secondary antibodies (1:500, AlexaFluor 568 Invitrogen) and mounted on SuperFrost Plus slides and coverslipped with ProLong Diamond Mountant (Invitrogen # P36970). Fluorescence images were acquired using an Olympus FV1000 confocal microscope. Antibodies were validated on positive tissues from CCI animals.

### Open field test (Supplementary Fig. 3)

The open field test consists of a dark Plexiglas enclosure (1 × 1 m) placed in a dark experimental room. Lighting was set as 16 lx in the open field. Rats were placed in middle of the enclosure and their exploratory behaviors were recorded with the ANY-Maze Tracking software for 10 min across the arena, virtually divided into 16 squares. The distance travelled and the maximal speed were measured during the first test minute for motor assessment. The time spent in the center of arena (four virtual central squares), was recorded over the 10 min test period for anxiety-like behavior assessment. The test was performed on day 1 and 4 of the injection regimen, for side effect assessment following acute and chronic drug exposure.

### WST-1 assay (Supplementary Fig. 3)

15,000 DRGF11 cells were seeded in quadruplicate in a 96-well plate in DMEM supplemented with 10% FBS, 2% penicillin/streptomycin and 25 mM Hepes. 24 h after cell seeding, 100 µl of fresh serum- and phenol red-free medium containing 10 µM of pepducin (0.4% DMSO), or corresponding scrambled control was added. After 24, 48 and 72 h drug incubation, 10 µl of WST-1 reagent (5 mg/ml) was added to each well and incubated for 4h at 37 °C, 5% CO_2_. The optical density (OD) of the formazan dye generated by the conversion of the WST-1 tetrazolium salt by mitochondrial enzymes was measured at 450 nm for each well using a GENios Pro plate reader (Tecan). Cell viability was measured using the following formula: (OD_450nm_ treatment/OD_450nm_ scrambled control) × 100.

